# Uncovering the translatome impact of transcriptome induced diversity in eukaryotes: framework and innovative insights

**DOI:** 10.1101/2024.03.08.584087

**Authors:** Paras Verma, Deeksha Thakur, Shashi Bhushan Pandit

## Abstract

Isoform diversity is known to enhance a gene’s functional repertoire. Insights into the extent of such sequence variability generated through alternative splicing (AS), may unveil the layers of gene function/regulation. Despite studies on transcriptome diversifying processes, the impact of AS or related processes on sequence diversity still needs to be explored. Current study presents an innovative framework that centralizes exonic loci while integrating protein sequence per entity with attention to splice site variability assessment. The resulting framework enables exon (features) to be tractable, facilitating a systematic, detailed analysis of isoform diversity. We analyzed isoform diversity in five representative organisms and detailed the role of AS and related processes influencing exon inclusion in imparting sequence variation for human genome. Through analyses of exonic variations in two maximally diverged isoforms of human genes, we unraveled intricate splicing patterns prevalent in coding and non-coding regions. We observed that alternative splice sites, sequence changes, and skipping of exons are prevalent in coding exons, while the alternate first exon events are predominant in non-coding exons. Our findings offer a comprehensive understanding of isoform diversity as a function of exonic entity framework, providing valuable insights into the orchestration of exonic events in shaping the proteogenomic landscape.

## Introduction

Gene architecture in eukaryotes facilitates generation of more than 1 mRNA per gene by composing differential combination of exons. This process of exonic region determination and subsequent combination is under regulation of several co-and post-transcriptional processes facilitating transcriptome (protein) diversification and helps define cellular level identities in higher eukaryotes (1–3). Consequently, dysregulation of this regulatory process could result in expression of aberrant transcript/isoform responsible for numerous pathological conditions (4). Among several mechanisms contributing to transcriptome variation, alternative splicing (AS) is widely studied, followed by alternative transcription initiation (ATRI) and termination (ATRT) processes. The latter gained substantial attention in the last decade with the advent of technological advancement in high-throughput mRNA sequencing (RNAseq) (5,6). Recently, AS/ATRI/ATRT processes have been unveiled to work simultaneously and provide feedback to transcription; for instance, AS-mediated splicing of introns significantly influences transcription rate, and this is noted to influence both co- and post-transcriptional splicing reciprocally (7–10). The collective influence is observed to affect both coding sequence (CDS) and untranslated region (UTR) regions from studies of the mammalian transcriptome. Their potential coupling (between the 3’/5’ UTR and the CDS region domains) has also been emphasized, implying their evolutionarily integrated role in enhancing overall diversity within the transcriptome (11).

The transcriptome diversity reflects a cumulative influence of ATRI/ATRT and AS processes. However, additional mechanisms exist in translation that contribute to shaping of proteome diversity. Alternate translation initiation (ATLI) and termination (ATLT) are widely acknowledged for diversifying the translation of mature RNA through the utilization of varied ribosomal entry sites, leaky scanning, re-initiation, or the inclusion of upstream ORFs in transcript structure (12–14). Some studies have also highlighted pervasive translation occurring from genomic regions (UTRs, introns, and long ncRNA), which traditionally exhibit lower specificity for ribosomal translation (15–18). Determining the impact of AS/ATRI/ATRT, ATLI/ATLT, or their combined interplay in generating proteome variation is challenging, which is further exacerbated by the realization that <30% of human exons protein-coding (19), and >4-fold fraction of alternative nucleotides (nt) participate in ATRI/ATRT than CDS region (20). Such complexity is not limited to humans but is also noted and found to be conserved in the mouse genome, where splicing exon profiles from non-coding regions were recapitulated for human chr21 (21). Notably, the exons involved in ATRI/ATRT processes and the ribosomal start/stop sites of ATLI/ATLT exons are predominantly conserved in orthologous genes across fungal, plant, insect, and mammalian species (22,23). Cumulatively, these findings raise several important questions regarding the mRNA-mediated effects of intron-exon definitions on resulting protein translation. Subsequently, it is intriguing to know how the coding region (CDS) accommodates alterations introduced by ATRI/ATRT and/or ATLI/ATLT and/or AS within intragenic transcripts. To the best of our knowledge, this unfilled gap in the understanding of translatome has yet to be addressed systematically in the context of uncovering splicing complexities.

Literature reports concerning splicing event elucidation relies majorly on pairwise comparison of transcripts. Over the years, numerous related methodologies were developed, that define and annotates the events, however encounters limitations in distinguishing their impact on CDS / UTR regimes. For instance, the numeric symbolic notation by Sammeth et al. (24,25) and utilization of splice graph-based data structures efficiently uncover similarity and complex events in transcriptome RNAseq data (26,27). Modifications of such associated approaches specifically for the splice graphs have been developed to uncover either the splicing complexity (27) with application in discovering unique exon junctions or to reduce the space and time complexity of associated calculations (28). These studies have emphasized the role of exons in activating alternate transcription initiation sites (7) and in hybrid exon quantification that may have the potential to modulate the principal protein sequence of the gene (29). However, these studies were limited to data derived from RNAseq and clearly lack the ability to identify other possible genes undergoing such processes and how proteins incorporate those changes. A systematic inferable integration of transcriptome and proteome data has been limited for reasons as reviewed in Reach-Solé and Years (30), wherein it has been emphasized that the ORF prediction lacks concordance with translation due to the difficulty in identifying choice of alternate translation sites. Notably, studies report consequences of such splicing induced dysregulation leading to several diseased conditions, where the expressed protein is from an alternate translation initiation sites instead of its cognate site (31). However, to decipher the nature of genome expression, it is essential to detect translation at the level of proteome. Moreover, delineating such events will also help uncover the resulting impact of ATRI/ATRT or/and ATLI/ATLT on AS.

To circumvent the challenges of integrating transcriptome with proteome, a thorough knowledge of the coding nucleotide coordinates of exons within CDS region, particularly in the context of AS, ATRI, and ATRT become imperative. For incorporating these and their associated aspects, we developed a systematic exon translational feature centric framework, Exon Nomenclature and Classification of Transcripts (ENACT) that streamlines computational tracking of exonic events, while facilitating straightforward interpretation. ENACT incorporates features for exonic loci in gene after comprehensively tracking their Coding Genomic coordinates (CGC) and genomic coordinates (GC) in isoforms as defined in RefSeq (32). The ENACT framework employs alphanumeric embedding alongside protein sequence features to unveil the extensive scope of diversity generated through transcriptome and proteome. Through ENACT and its assisted analysis of isoforms coded per gene, we showed that distinct splice site variations and translational features are associated in imparting sequence space differences from reference isoform. Current analysis unraveled association of exon skipping and alternate splice site choice in imparting sequence diversity. We studied transcript model annotations harboring such events as case to showcase their possible roles in imparted isoform diversity. Our work emphasized that even though overall fraction of A(ss) events to be preponderant AS type has declined in animals, their distinct role can be noticed from their positional prevalence in gene architecture that may be facilitating mechanism to alternate translation initiation and termination. Additionally, to improve current understanding of AS impact and for interest of scientific community, we made ENACT representations publicly available as ENACTdb (http://www.iscbglab.in/enactdb) database. The information in it is integrated with additional attributes of predicted Pfam domains and secondary structures with isoform represented through an interactive display of features employing state of the art web-design tools as ReactJS and Django.

## Materials and methods

### Overview of ENACT framework

#### A) Unique indexing of exon to construct gene architecture and define alternate / constitutive feature to exon

For a given gene, we select an isoform having the maximum number of coding exons from a set of curated isoforms (RefSeq proteins having ‘NP_’ prefix) and define it as a Reference ISOform (RISO). If the number of coding exons is the same in two or more isoforms, then the one with the longest length is selected as RISO. If a gene has no ‘NP_’ prefixed isoforms, then RISO can be chosen from all known isoforms using the similar criterion (Box –I).

Exons of RISO, primarily, constitute the Reference Set Of Exons (*RSOEx*), which are later populated with exons from other isoforms (*nrIsfSet*) based on GC overlap (Box-I and represented in Figure 2). This procedure of exon selection (for *RSOEx*) involves segregation of overlapping exons (*OlEx*, with GC overlap to *RSOEx*) from non-overlapping exons (*NolEx*, without GC overlap to *RSOEx*). *NolEx*, could have individual candidates (*NolEx-A*) which are added to *RSOEx* and subgroup of exons (*NolEx-B*) with self GC overlaps (routine *defineSuboverlapExons* (Box-1), Figure 2B1). Subsequently, *NolEx-B* subgroups are processed to identify *Qualifier_exon_* in following way: from a subgroup (*NolEx-B*), an exon of minimum length of at least ten amino acids (30 nt) or the longest (if maximal exon length among overlapping entities (*GoEx*) is <30 nt), is chosen as representative exon (*Qualifier_exon_*) for *RSOEx* set (Figure 2B1). Others left in the above subgroup are moved to *Exon_variant_*.

Resulting *RSOEx* are numerically sorted considering their GCs, and their linear index (position in gene) is depicted as the first character in Block-II. The next character of Block-II (Figure 2E) depicts the alternate/constitutive nature of exons. Defined by GC consistency, an exon is considered alternate (shown as ‘A’), if it lacks uniform presence in all transcripts and constitutive (depicted using ‘G’) otherwise. Additionally, certain exonic positions are depicted using letter ‘F’, representing them being present in all isoforms, however, with splice site variations.

#### B) Relationship definition between *RSOEx* and *Exon_variant_*

The previous step identifies reference exon (*RSOEx*) and variant exon sets (*Exon_variant_*). In this step, we proceed to define specific splice site variant relation (5’ and/or 3’ exon) considering their GC overlap. We chose to prefer exon definition model and focused on exonic entity while considering affected splice site (33), as ENACT framework involves their featurization with protein sequence. Each exonic entity in *Exon_variant_* set is compared with those in *RSOEx*, taking into account their GC overlap, and are assigned following notation depicting splice site relation:

- n: It denotes different 5’ splice site (5ss) but identical 3’ splice site (3ss) of an *i^th^* entity of *Exon_variant_* to the *k^th^* entity of *RSOEx*.
- c: It denotes identical 5ss but different 3ss of an *i^th^* entity of *Exon_variant_* to *k^th^* entity of *RSOEx*.
- b: It denotes different 5ss and 3ss of an *i^th^* entity of *Exon_variant_* to *k^th^* entity of *RSOEx*.
- 0: It denotes identical 5ss and 3ss of *i^th^* entity of *Exon_variant_* to *k^th^* entity of *RSOEx*.

The above notations (n/c/b/0) are utilized as first character of Block-III and represents (for n/c/b) either extension or shortening of the exon length. It should be noted that, when exonic entity *ith* of *Exon_variant_* showed GC overlap to more than one entity in *RSOEx*, we dealt those cases separately as intron retention events.

### Occurrence of splice site changes

The variant relationship between entities of *Exon_variant_* between *RSOEx* are facilitated by their GC overlap. To accommodate and acknowledge more than 1 GC overlapped entities in *Exon_variant_* with that of single entity in *RSOEx*, we track such instances by counting their occurrences. For instance, *i^th^* and *j^th^* entities of *Exon_variant_* show identical 3ss but different 5ss (n variation) with *k^th^* entity of *RSOEx*, here we track their unique occurrences i.e. differing 5ss in *i^th^* and *j^th^*exons. It is represented as a numeric character in the 2^nd^ position of the Block-III (Figure 2). The default value of the character is 0, which incremented by 1 whenever unique (not observed before) splice site is encountered. Interestingly, the number of n/c/b events of a specific exon locus (determined by Block-I linear indexed attribute) in a gene can be obtained by looking at this position.

#### C) Amino acid coding (translational) attribute of exon those defined in *RSOEx* and *Exon_variant_*

As described above, exons listed in *RSOEx* and *Exon_variant_* are characterized with attributes of position, prevalence and nature of splice site variation (n/c/b/0) (Figure 2B and 2C). As exon can be a part of noncoding (UTR) or coding region (CDS) in isoform, we incorporated attribute unveiling its scope and contribution to isoform sequence. We used two characters to describe the coding status attribute in the Block-I (Figure 2E). The first character denotes amino acid contribution to isoform and the second character identifies the general nature of an exon such as coding/noncoding or both. The first character uses following numeric notation for its isoform specific scope:

- -2: It denotes that the exon contributes no amino acids.
- 0: It is a primary placeholder to depict the M case (single nucleotide exons)
- -1: It depicts premature stop codon in the upstream exon; hence, this exon does not contribute to the amino acid sequence, even though it has more than one nucleotide in the CGC.
- 1: It shows that the exon contributes amino acids to the isoform.
- ≥2: It is a counter denoting the number of different amino acid sequence variants observed for an exon with the same GCs.

The above code depicts the isoform specific scope informing exon behavior in its harboring isoform. To unveil translational behavior throughout in gene architecture, we use the following alphabet notation:

- T: It depicts that an exon (or its splice site variants) always codes for amino acid sequence or is a part of coding sequence (CDS) in a transcript. The exon region has a defined Coding Genomic Coordinate (CGC) in all transcripts.
- U: It shows that an exon (or its splice site variants) forms the untranslated region of transcripts whenever it occurs or does not have CGC.
- D: It shows that an exon (or its splice site variants) is part of CDS in at least one transcript and is part of UTR in another transcript(s). Notably, these exons should share identical Genomic Coordinates (GC) across transcripts existing as either coding or non-coding.
- M: It is assigned for single nucleotide protein coding exon in the CGC region.
- R: It is for an intron retention exon, described later in the nomenclature section.

The characters are joined by ‘.’ (in the order of Block-I, Block-II and Block-III) as demonstrated in Figure 2E, depicting a 6-character long alphanumeric string descriptor, which we term as Exon Unique Identifier (EUID).

### Intron Retention (IR)

It is a special case of exon nomenclature where the EUID is insufficient to capture details of an intron retention event (as it involves retention of intron region between exons). To describe this, we used EUIDs of two exons and merge them in five separate identifiers combined with the ‘:’ colon symbol to define IR cases. The first identifier is the alphabet ‘R’ to recognize that the exon is involved in IR, followed by a digit describing its amino acid coding attribute (the same notation is used as described before). The third and fifth identifiers are exon EUIDs, between which the intron/exon region is retained to form the IR exon. The fourth identifier is a numeric character showing the number of retention events observed involving exons and their variants. We use 0 as the default value of this counter.

R:1:U.-2.A.2.n.1:0:T.1.A.3.0.0

The above IR exon depicts that it is an amino acid coding exon having the retention of intron region between exons U.-2.A.**2**.n.1 to T.1.A.**3**.0.0. It is the first instance involving exons 2 and 3 (shown in bold as these are the relative positions of exons in a gene). The same is illustrated in Figure 2D2.

### Overview of dataset collection, defining mIDI and sequence identity/coverage

The isoform/transcript sequence and other accessory information (Genomic coordinate (GC) and Coding genomic coordinate (CGC)) of genes encoded in five representative organisms were obtained from RefSeq database of NCBI (34). We computed isoform diversity for a gene set (GeneISF2) consisting of genes with at least two distinct isoforms, which could vary in their amino acid (’aa’) lengths or/and sequence. The GC/GCG was obtained for each gene in the gene table format was downloaded from NCBI. For each gene, a maximally divergent isoform is defined as the one having minimum sID to the Reference Isoform (RISO).

To compute sID and sCOV, we performed local pairwise sequence alignment using *Align* routine of Bio.Align module of Biopython (35) package. The default parameter for alignment used was BLOSUM62 matrix, gap open and extension penalty of -10 and -0.5 respectively. The sID is computed as (number of identical residues/alignment length). The sCOV is computed as (number of aligned residues of RISO/length of RISO).

### Event definitions for alternate translation/transcription initiation/termination

For the genes participating in the given sID and sCOV bins, following events are evaluated.

#### Alternate first and last exons

The alternate first and last exon corresponds in mIDI isoform to as different initial and terminal exons. For the coding instances, 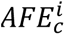 and 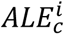 are evaluated to true for gene ‘i’, if index of their first coding exonic loci and last coding exonic loci instance are not identical in mIDI and RISO ( see 3, 4 and 5)

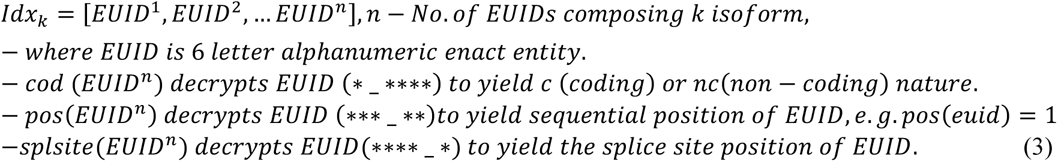

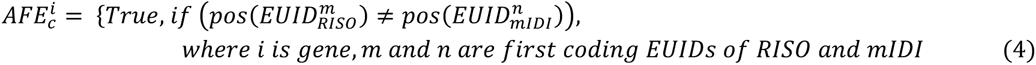

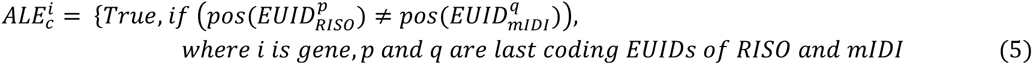

Their definition of their non-coding instances (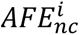 and 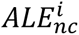) evaluates to true for gene i, provided either of *mIDI* or *RISO* have UTR exon loci instance in those region (see 5 and 6). If exon loci instances are available in only *mIDI* but not in *RISO* or vice versa for 5’ or 3’ UTR regions, condition is evaluated to be true. If either region (5’ or 3’ UTR) has exonic loci for both *mIDI* and *RISO*, then match in exonic first and last exonic loci for those regions are evaluated.

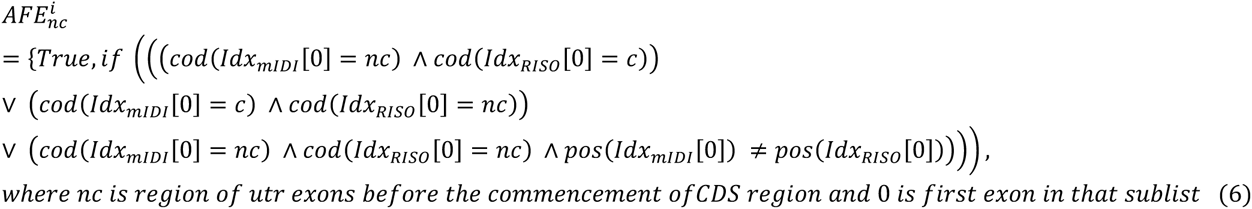

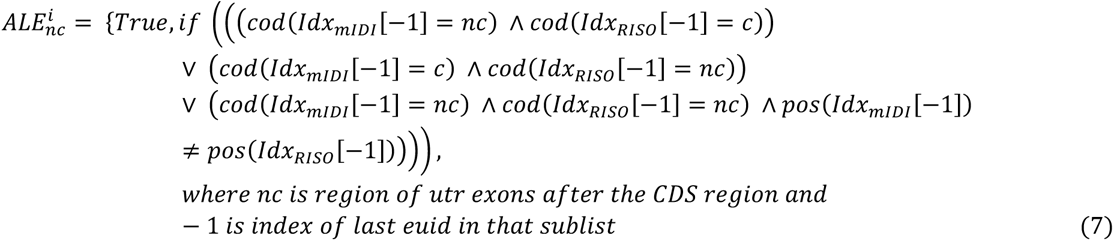

#### Exon gain and skipping events

Identified ES (lost exon to RISO) and EG (gained exon to RISO) events were made distinct from loci which can be candidates of AFE and ALE. For the non-coding counterpart following conditions are implied, and skipped exon is considered such that exon loci (Block II) that are present in RISO but not in mIDI are bound to occur within first and last exonic loci of theirs (see 8). Similar evaluation is considered for EG; however, exon set is reformulated for loci that are present in mIDI but not in RISO (see 9).

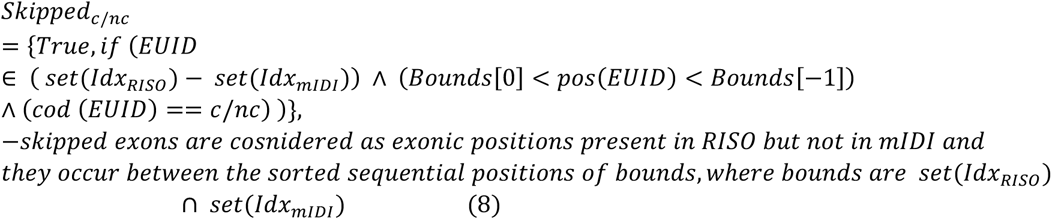

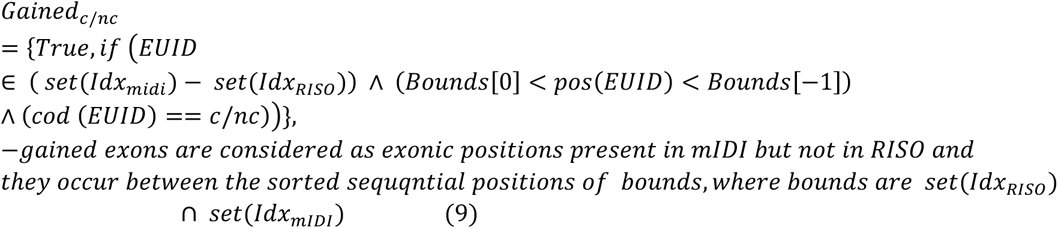

#### Alternate splice site choice exons

A(ss) events are evaluated true for gene i, provided there exists an exonic loci in mIDI, identical to RISO (indexed position), however but with splice site variation to RISO entity (see 3 and 10).

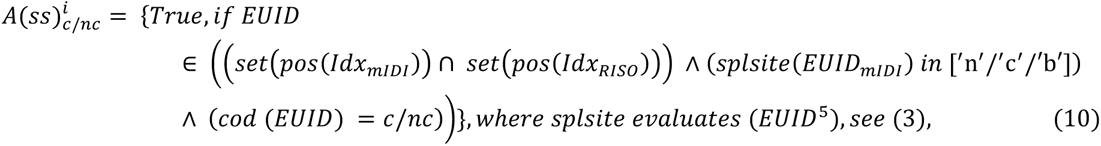

#### Change in sequence events

Exonic entities with T and D tag in the Block-II (pseudo-global scope) can undergo variable isoform specific scope which differs from the *RSOEx* instance. These events are evaluated to be true when RISO and mIDI shares identical Block-II linear index and variable local status. Their depiction by Sv1 and Sv2, is defined as follows:

Sv1: exon pairs (mIDI and RISO instance), that are subset of one another, while showing length change ≥ 2aa and overhanging to one or another splice site start/end (see 11).
Sv2: exon pairs (mIDI and RISO instance), that are >2 aa length differing but not showing splice site overhang or exon pairs that are of comparable length (<= 2aa) but not having subset of shared sequence (see 12).

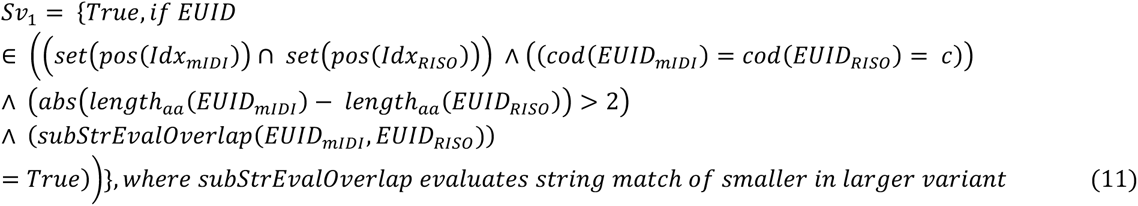

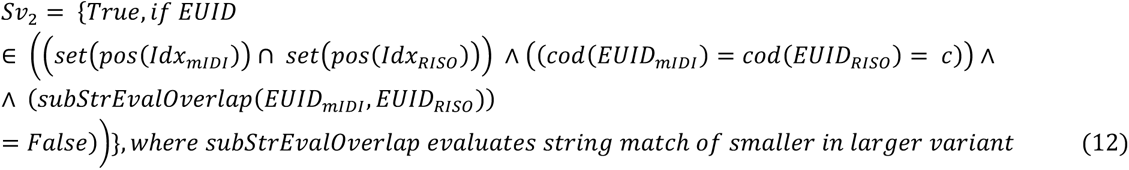

## RESULTS AND DISCUSSION

### Maximal sequence diversity is observed in human and mouse isoforms

To assess isoform sequence diversity generated due to a combination of AS and other transcriptional/translational processes, we relied on accurate GC and CGC available in the NCBI’s RefSeq transcript models. We defined each gene’s maximal isoform sequence diversity by recording the minimum sequence Identity (sID) between isoforms to the Reference ISOform (RISO), terming the other isoform as mIDI (see methods). We characterized the isoform diversity by the sID between mIDI and RISO and their alignment coverage (sCOV) (see methods) normalized by RISO length. A high sID and low sCOV would indicate only a local high similarity between two isoforms, whereas a low sID with low sCOV indicates poor similarity between them.

To explore isoform diversity, we examined the joint distribution of mIDI’s sID and sCOV alignment parameters from RISO concerning gene set GeneISF2 (see methods) in five representative genomes (see figure 1 and table 1). In general, sID and sCOV are high across genomes, and the same is evident in Figure 1, which shows most genes (∼98%) have sID >50%. However, these have varied sCOV (maximal variation observed in *C. elegans* and *H. sapiens*), indicating that only partial region of mIDI sequence is aligned with RISO. *C. elegans* showed maximal number of genes (∼23%) having mIDI with sCOV <50%, followed by human where this fraction is ∼11%). Moreover, genes in these genomes exhibit the maximum difference between RISO and mIDI with mean difference length of 159, 124.1, 105.8, 76.9, 69.3 aa for *C. elegans*, *H. sapiens*, *M. musculus*, *D. melanogaster*, and *D. rerio*, respectively (Supplementary file SuppDataExcel.xlsx, Worksheet: F1_Identity_Coverage_gene_wise). A small fraction of genes (<2%) show diverse sequences with varying sID’s and sCOV. Considering both of these parameters, we observed that genes having isoforms with high sID (>50%) and low sCOV (<50%) are more prevalent in worm (20%), followed by human (9%), mouse (6%) fruit-fly (5%), and zebra fish (3%) genomes.

**Figure 1:**
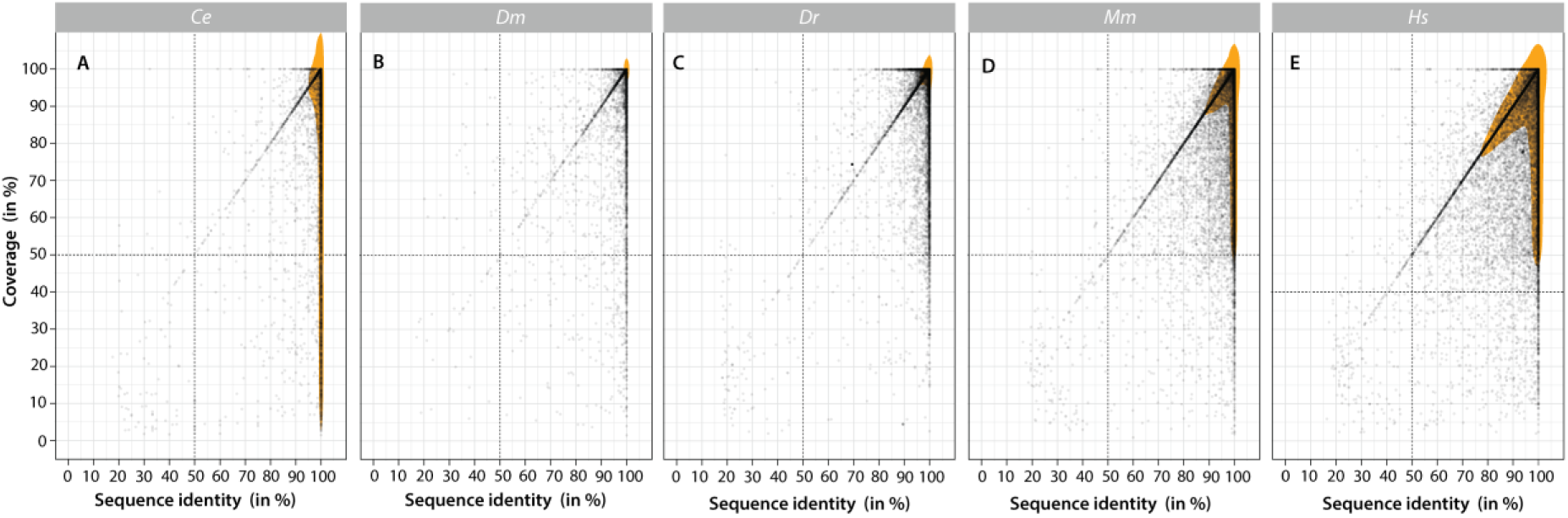
Isoform diversity comparison of protein coding genes encoded in genomes of five model organisms. The pairwise sID and sCOV between RISO and mIDI isoforms are shown as scatter plot for genes from *C. elegans (Ce)*, *D. melanogaster* (Dm) , *D. rerio* (Dr) , *M. musculus* (Mm), and *H. sapiens* (Hm). Each dot represents a gene in the plot. The transparent orange shared region represents the 2D kernel density estimation (using function stat_density_2d, ggplot 3.3.0 in R) of distribution of genes in the plot.

**Figure 2:**
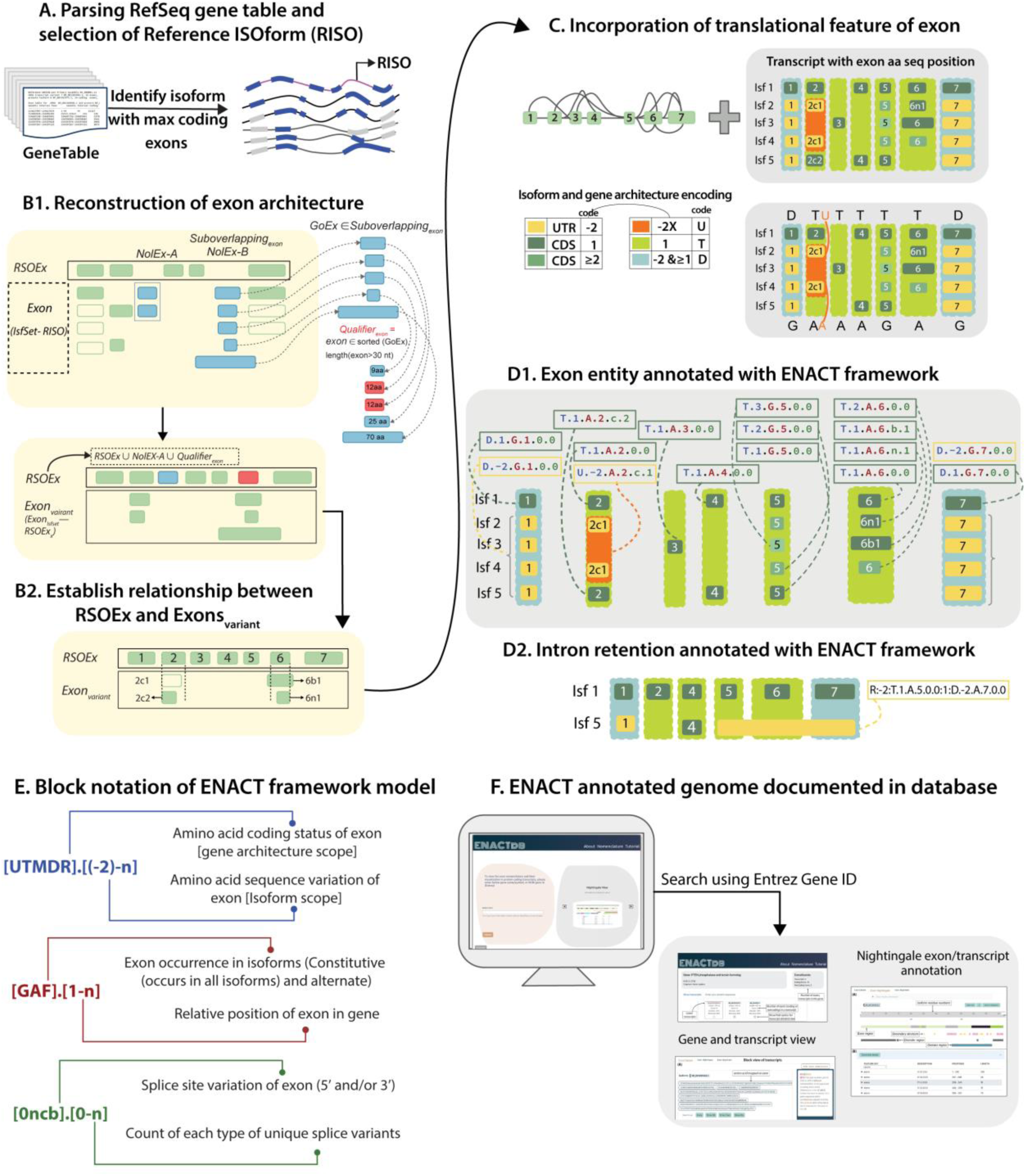
Overview of Exon Nomenclature and Classification of Transcripts (ENACT) framework. The main steps of the ENACT algorithm (A-D) and the framework model (E) are illustrated for a gene. A) The source data of genes, comprising curated models of transcripts with GC and CGC concerning each exon is obtained from RefSeq in gene table format (Blue and grey color regions represent coding and non-coding exons, respectively). The reference isoform (RISO) is selected based on the maximum number of coding exons (see methods). B1) RISO exons constitute primary set of reference exons (*RSOEx*), which are later screened for candidate exons from other isoform: a. singleton non-overlapping (concerning GC of *RSOEx*) exons (*NolEx-A*) and b. a selected exon (Qualifier) from a group of overlapping (*NolEx-B*) exonic entities (based on their GCs) that has a minimum length ≥30 nt or the maximum length. Thus, *RSOEx* set consists of non-overlapping exons (sequentially ordered based on GCs) from all isoforms. A set of Exon_variant_ is maintained comprising entities with GC overlap to *RSOEx*. An exon with GC overlap with ≥2 *RSOEx* instances is tagged as Intron retention and maintained separately. B2) Exons in Exon_variant_ are compared with those of *RSOEx* to define splice site variants (alternate splice site) as n/c/b instances (see methods for details). C) CGCs are used to establish translational attributes for each exon (*RSOEx*/*Exon_variant_*). The relative position (considering their GCs) of all exonic entities (*RSOEx* and *Exon_variant_*) and their prevalence in isoforms are noted as linear index and constitutive/alternate state attributes. Thus, each exon is assigned its relative position, translational feature, occurrence, and splice site variation (if known). D1) The exon attributes noted in the previous steps are combined to construct a 6-character alphanumeric notation defined as exonic entity (EUID). D2). The intron retention instances are identified in step B and annotated with IR codes. E) The 6-character EUID is segregated into three Blocks (I, II, III) of 2 characters each. G) The description and notation for each exon character is illustrated in the figure. The annotated exons of genes encoded in five model organisms are documented in (ENACTdb) database with distinct visualization views of exon annotations.

**Table 1:** Summary statistics of sequence sID and sCOV between RISO and mIDI.

| Organism | No. of genes in GeneISF2 | <sID> (sd) | <sCOV> (sd) |
| --- | --- | --- | --- |
| *C. elegans (Ce)* | 4467 | 95.6 (11.2) | 72.2 (27.6) |
| *D. melanogaster (Dm)* | 7053 | 97.0 (8.8) | 90.8 (17.6) |
| *D. rerio (Dr)* | 9810 | 95.9 (9.8) | 89.4 (16.7) |
| *M. musculus (Mm)* | 13749 | 94.1 (11.0) | 81.7 (19.6) |
| *H. sapiens (Hm)* | 14761 | 90.9 (13.2) | 78.2 (20.8) |

Given the relatively higher proportion of isoform variation in human, we focused our attention towards them for a detailed investigation of AS processes and their contribution to sequence diversity. Although associating protein sequences to exons for elucidation of splicing events seems trivial for a single transcript, it is challenging to compare exonic entities across multiple transcripts, especially with the possibility of varying CGC and GC of exon loci. The challenge becomes further compounded when other transcript variations arise from ATRI/ATRT and ATLI/ATLT events, which could result in similar/variable translational initiation and stop sites (TIS/TSS) while simultaneously having splice site changes in exons. To comprehensively address these issues, we formulated an innovative ENACT framework to uniquely annotate exons for their attributes, such as coding ability, relative position, and splice site variations.

### Incorporating exon splice site and sequence variations unveil exon multifaceted features (ENACT framework)

To emphasize variability embedded in the exon translational attribute, which could be either local (transcript specific) or global (gene architectural), we have focused on establishing linear indexing of exon first followed by characterizing splice site variability. Figure 2 shows the main steps in exon annotation under ENACT framework. Below, we describe the following main steps:

#### a. Maximizing exon coverage of gene architecture

The approach for linear indexing of exons and maximizing their coverage is described in Figure 2A. We select a Reference ISOform (RISO) with the maximum number of coding exons among curated isoforms. The exons of RISO constitute the Reference Set Of Exon (*RSOEx*), which is populated with intervening (non-overlapping) exons obtained from other transcripts. This step involves identifying exon(s) (one or more) from other transcripts that lack GC overlap with the reference isoform (RISO), referred to as Non-overlapping exons (*NolEx)*. Among *NolEx*, exons having identical GC (*NolEx-A*) in multiple transcripts are appended to *RSOEx* (Figure 2A). The rest of the exons of the *NolEx* set, which lack uniform GC across their occurrences in transcripts, indicating their splice site variability are referred as *NolEx-B* set (Figure 2A) and dealt in groups of overlapping entities. To screen candidate exons among all possible sets of overlapping *NolEx-B* exon sets (exon GC’s will be overlapping intra and non-overlapping to inter *NolEx-B* sub sets), we optimize exon selection through an iterative procedure, where we select the exon with the minimum length of at least 30 nucleotides as representative for *RSOEx* (*Qualifier_exon_* of *NolEx-B* in Figure 2A and *Suboverlaping_exons_* in methods). The qualifier exons are added to *RSOEx* and they are indexed linearly based on their genomic coordinates. The *RSOEx* exons represent a comprehensive collection of non-overlapping exons while other exons (*Exon_variant_*) are *RSOEx* splice site variants. Resulting *RSOEx* exons have unique relative positions, constituting one attribute under ENACT framework.

#### b. Splice site relationships between *Exon_variant_* and *RSOEx*

The splice site variations of an exon are defined based on GC overlap between *Exon_variant_* and *RSOEx*. We examined exon instances overlapping with only a single *RSOEx* entity for their GC coordinates (5’ and 3’ exon splice sites) to denote their type of splice site variations (n/c/b) compared to *RSOEx* (Figure 2B). The n, c, and b characters depict alternate 5’ (with unchanged 3’), alternate 3’ (with unchanged 5’), and both altered splice sites, respectively (see methods). Additionally, for every instance of unique splice variation for an *RSOEx* entity, we maintain a specific count record. These two characters adds up to ENACT exon attributes. Importantly, we filtered exon instances in the *Exon_variant_* set overlapping with two or more exons in *RSOEx* (or their splice site variations) as a presumptive set of intron retention exons (IR events).

After indexing the relative position of reference exons and establishing Exon_variant_ as their splice site relatives, we associate coding status (amino acids contribution to isoforms) to exons considering their CGC (translational feature). An exon entity and its variant could have the same GC but variable CGCs in two or more transcripts, giving rise to variable amino acid (aa) sequences from the same exonic loci. Additionally, absence of CGC can lead to exon being coding in some isoform(s) and non-coding in others. For instance, as illustrated in Figures 2C/2D, the exonic loci 1 and 7 are coding in ISF 1 and non-coding in all other ISFs. Exonic loci 2 of *RSOEx* and one of its splice variants are coding in ISF 1 and ISF5, respectively, while another variant (2c1) is non-coding in ISF 2 and 4 (Figures 2C and 2D). In order to delineate and distinguish such events, it is essential to acknowledge that the scope of coding status extends globally (gene architecture) and locally (transcript instances). Under ENACT framework, two distinct characters represent these exon features: one indicating local coding status and the other representing pseudo-global coding status (see methods).

In Figure 2C/2E, variations in amino acid coding status are illustrated through color-coded exons to facilitate easy distinction. As depicted in fig 2C/2E, the exon-2 variant (2c1) in ISF2 and ISF4 is non-coding, so it has a pseudo-global scope of U and ’-2’ as local status. However, the parent form of exon-2 is amino acid coding wherever it occurs and is assigned ’T’ and ’1’ as pseudo-global and local characters, respectively. The exon loci positions 3, 4, 5, and 6 are coding in all isoforms, so these are assigned ’T’ as pseudo-global code. However, it is pertinent to note that exon-5 exhibits amino acid sequence variation (represented through shades of green color) without splice site change, possibly due to varying TIS in ISF 2, 3, and 4. These exon variation instances are shown by changing local transcript characters as 1, 2, and 3 under the pseudo-global ’T’ tag. A similar scenario is observed for exon-6, where an amino acid change is evident in ISF4. The splice site variants (6n1 and 6b1) of exon-6 have local coding status of ’1’ because these are separate splice variant instances compared to *RSOEx*. The exonic loci 1 and 7 have ’D’ assigned as gene architecture scope as these are coding in ISF1 and non-coding in the other isoforms (Isf-2 to Isf-5).

The translational potential of an exon captured at local, isoform-specific information, and global at gene architecture level is merged with exon relative indexed position and splice site variant notation, as described in steps (a) and (b). Combined characters describe attributes of Exons and composes its Unique Identifier (EUID). Once formulated, a subset of EUID characters can be used to construct sub features by invoking EUID^k^, where ’k’ could represent one or more attribute or their combination. We split EUID into three blocks, each encapsulating specific characteristic features of exons. The first block (blue-colored characters in Figures 2D and 2E) denotes the coding nature of exon instances, wherein the initial character depicts the gene architecture scope (pseudo-global), and second character enlists local isoform-specific information. The second block (dark red-colored characters in Figure 2D and 2E) indexes the relative linear position of the exon in a gene (step a), followed by a feature of exon occurrence as alternative (in some isoforms) or constitutive (all isoforms) denoted by characters A and G, respectively (illustrated in Figure 2C). The final Block (dark green-colored) depicts the exon splice site variations encoded as n/c/b and its unique instances. A special notation of 0.0 is assigned to exons in *RSOEx*. Notably, Blocks-II and III establish a detailed relationship between exons across transcripts, enlisting their all possible variations while incorporating any combination for Block-I (illustrated in Figure 2C). The intron retention is filtered in step (b), which is defined as a combination of annotated exons with their EUIDs and additional information on their coding/non-coding nature (Figure 2D) (see methods).

We annotated exons using ENACT framework for the five model organisms. Their exon featurization and transcript annotation have been documented in ENACTdb (Figure 2F and Supplementary section S3). Additionally, for each exon, various predicted features are superposed for enhanced visual representation, serving as necessary aid to comprehend the complexities presented in gene architecture. Below, we discuss utility of ENACT alphanumeric exon annotation to delineate AS events to understand their impact on isoform sequence variations.

### Analysis of the contributions exonic variation(s) on isoform diversity

We assessed the isoform sequence diversity of genes through the maximal variation in sID/sCOV between mIDI and RISO isoforms (Figure 1). This diversity could arise from combining one or more exonic variations (such as n/c/b, amino acid sequence changes) or exon inclusion/exclusion in a transcript/isoform. To uncover the extent of various exonic variations and their contribution to isoform diversity, we analyzed differences between the exon features of the mIDI and RISO isoforms. As this would require comparisons of shared exonic variations, we employed the ENACT framework, which provides the relative position of exons with their splice site or/and sequence changes attributes. The discretization of features in ENACT enables quantitative comparisons of exon features, such as amino acid sequence variations (Block-I), relative position (Block-II), and splice site variants (n/c/b) of Block-III (Figure 2D). It is pertinent to note that comparisons of Block-I and III features would identify exon variations, and Block-II provides information on inclusion/exclusion between isoforms. Using ENACT annotated exons, the difference in the number of exons (*δ_exon_*) between mIDI and RISO isoforms is compared to the fraction of shared exons having identical features (*FrSim_exon_*) for decoding exonic variations responsible for sequence diversity. The *δ_exon_* and *FrSim_exon_* are given by equations 1 and 2, respectively:

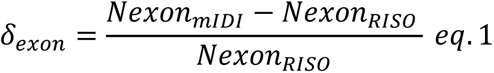

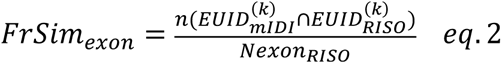

where *Nexon_mIDI/RISO_* refers to number of exons in mIDI or RISO isoform. The *δ_exon_* of 0 would correspond to same number of exons between mIDI and RISO. Whereas, a positive and negative value would refer to a relative greater number of exons in mIDI and RISO respectively. A 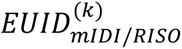 is a set of exons described by their *k* positional characters from mIDI or RISO, for example 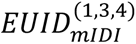 would refer to set of EUID elucidates exon features from mIDI described by 1^st^ (amino acid sequence status), 3^rd^ (relative position of exon) and 4^th^ (type of splice site variant) attributes. The numerator in *eq.2* refers to the number of exons sharing similarities for features in this set, later normalized by exon count in RISO. A value of 0 refers that no exons of mIDI and RISO share any feature (subset characters of EUID) and a value of 1 would represent that shared exons of mIDI and RISO are identical as in RISO with features. The maximum possible FrSim_exon_ is given as ratio of number of exons in mIDI to RISO (see Supplementary information S1).

We analyzed exonic variations of human genes and grouped them in sID bins of <50)%, 50-75)%, 75-90)%, 90-100)%, and 100% to explore the preponderance of types of exon variation with isoform divergence. For each bin, we assessed the shared exons attribute(s) similarity (*FrSim_exon_*) at decreasing levels of stringency for matching exon attributes. Initially, we considered exons having an identical amino acid sequences, splice sites, and relative positions (Figure 3A), followed by identical amino acid sequence and relative positions (Figure 3B), and lastly, the same relative positions (Figure 3C). The last condition will provide the extent of the maximal match possible between mIDI and RISO, while also revealing exon skipping AS event. Meanwhile, the first two former levels of exon match provide their variations. The decreasing level of stringency of shared exons is depicted in the vertical panels of Figure 3, whereas the horizontal panel exhibits them with increasing sID. Figure 3A shows small number of genes (∼2%) have high sequence divergence (<50% sID), and most (∼69%) have low divergence (≥90% sID). To quantitatively analyze the distribution pattern of gene density between δ_exon_ and exonic features, we concentrated on two regions: one represents genes having lost up to half of their exons while preserving at least half of their shared exon features (B-region, shown in the blue colored box of Figure 3) and the other region (D-region, shown in a green-colored box of Figure 3) is a subset of B-region, showing maximum exon feature similarity between mIDI to RISO. As seen in Figure 3A1, genes having high divergence are widely distributed, and ∼24% (B-region) of these shows >50% of their similarity in exonic features (sequence, position, and splice variants), despite losing half of their exons relative to RISO. Among these, genes with maximal *FrSim_exon_* scores are observed only for a small fraction (0.4%, along D-region), which have identical exon features in both mIDI and RISO. The relative fraction of genes in the B-region or D-region increased with sID (Figure 3A), except for 100% sID bin, indicating genes with low diversity tend to maintain similar exon composition and identical exonic features. Interestingly, genes with 100% sID isoforms show a wide distribution of exon attribute similarity for the same number of exons in mIDI and RISO (Figure 3A5). At high sID, we also observed that a number of genes have more exons in mIDI compared to RISO (δ_exon_ >0, Figure 3). It is pertinent to note that exon count includes both UTR and CDS exons, so the *δ_exon_* could be the same, while the number of shared coding exons could vary. This observation could also be due to the partial alignment of isoforms, which is apparent from their average sCOV. Surprisingly, we also observe several genes consisting of isoforms with no similarity in shared exons (*FrSim_exon_*=0) across sequence identities. For instance, gene crystallin gamma N isoforms NP_653328.1 and NP_001295221.1 share 68.5 sID has no identical exons considering all three features.

**Figure 3:**
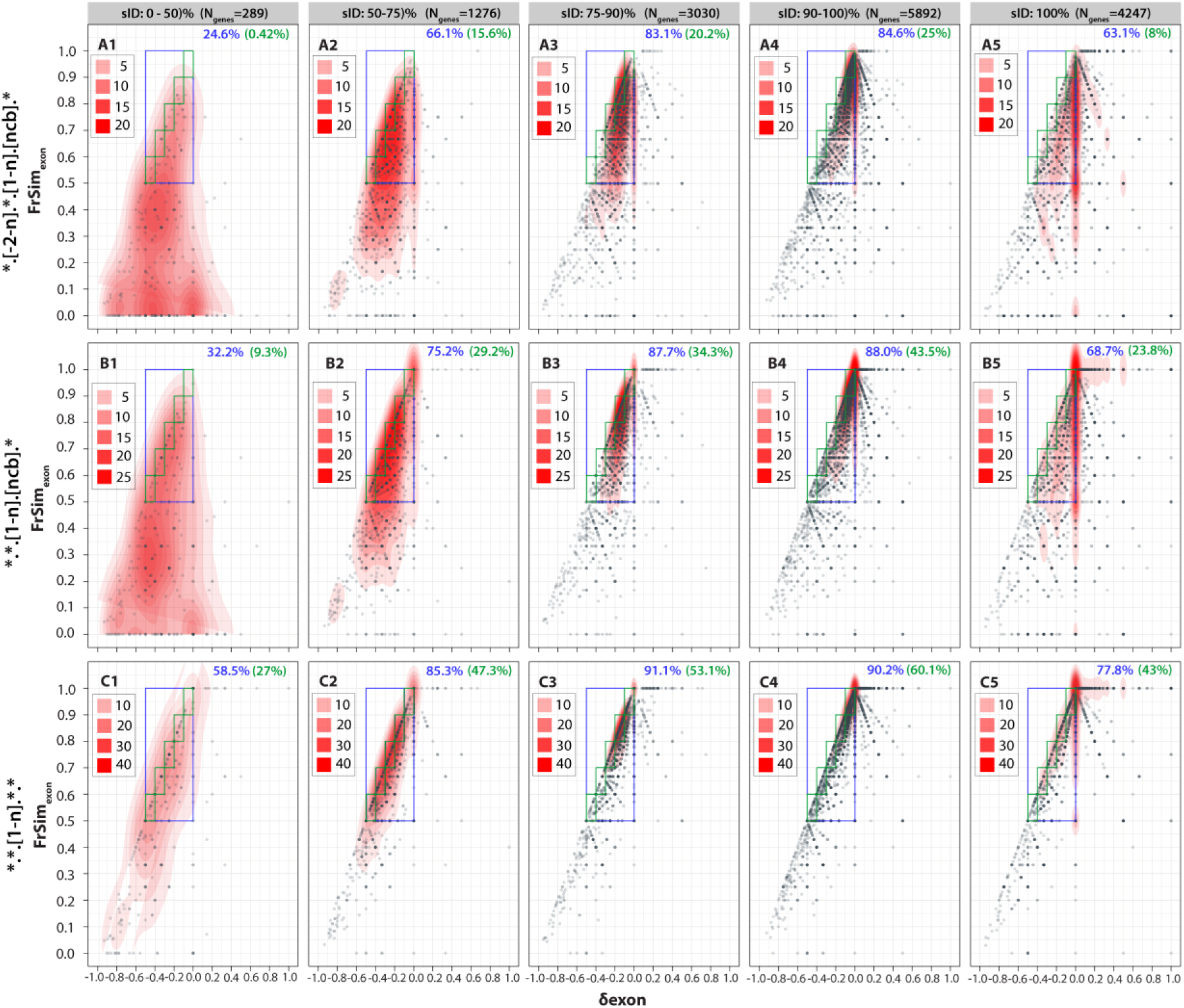
Differences in exon composition and feature variation between mIDI and RISO across various sequence identities. The comparison of exon feature similarity with exon variance between mIDI and RISO (of a gene) is shown as a scatter plot for various sequence identities representing the isoform diversity of a gene. The number exon variance between mIDI and RISO is compared with shared exon features, representing varying stringency as the number of matched features (shown in vertical panels). This comparison is performed for genes grouped based on sID bins (shown in horizontal panels). Each dot in the scatter plot represents a gene with the X-axis as δ-exon and Y-axis as *FrSim_exon_* similarity, with both measures being normalized by RISO exon counts (27 genes had δ-exon value >1 and were not included in this analysis). The measure of the similar exon is quantified by: A) considering all Blocks (I: translational feature, II: linear exons loci indexing, III: splice site variation), B) considering only Block-II and Block-III, while C) considering only Block-II (exons relative position). The red color shaded region represents 2-D kernel density estimation (using function stat_density_2d under ggplot 3.3.0 in R). The blue-colored box shows region of plot having mIDI having lost up to 50% of their exons and containing more than 50% of exon feature similarity. The green colored region envelops the box of 0.1 width (as diagonal region) of the blue box. The percentage of genes observed in the blue and green colored boxes are mentioned in each figure panel.

Upon relaxing stringency of matching exon attributes to compute their similarity (Figure 3C), we observed an increase of gene fraction in B-region/D-region as well as a decrease in genes with no shared exonic features (*FrSim_exon_*=0) across various sID bins. The drastic increase in exons having identical features indicates that most genes have mIDI varying from RISO predominantly in their amino acid sequence feature. Moreover, a significant fraction of genes in the B-region lie along the maximal similarity matches of exons (Figure 3B) that is given as the ratio of mIDI to RISO (Supplementary information S1). Finally, we examined the distribution of genes when only relative exon positions are identical in both mIDI and RISO (Figure 3C). As mentioned before, it can infer exon skipping AS events, thus providing insights into sequence variation generated due to the presence of exons from another locus. It is evident from Figure 3C that gene fraction along maximal exon match (D-region) increases drastically across sequence identities. For highly diverging isoform containing genes (sID <50%), ∼58% have at least half of their mIDI exons loci shared with RISO, suggesting variation in most genes is probably driven by coding exons from other exonic loci. On the contrary, low-diverging isoforms have most genes (>77%), showing more than half of shared exons, indicating a minor contribution to isoform divergence. Surprisingly, there is no noticeable effect of relaxing Block-I matching criterion (relative position) in 100% sID bin, emphasizing that most isoform variation is primarily due to splice site variations of exons. With increasing sID, we noted dissimilar roles of splice site choice and translation exon features. Isoforms with low divergence (>90% sID) would likely have similar underlying shared exons. However, we have observed noticeable increase in the gene fraction (Figure 3C5) for the B-region, unlike in highly diverged isoforms. We also observed modest gene population having identical numbers of exons in mIDI and RISO, while showing dissimilarity in *FrSim_exon_*. Additionally, we observed higher gene density in region of *FrSim_exon_*=1 when exon relative position is considered (Figure 3C). It is important to recognize lower fraction of matched exons for considerable gene count can also be a consequence of poor sCOV.

Above analysis clearly indicates preferential role of splice site variations and translational feature of exons leading to isoform divergence across sID bins, suggesting potential role of them and their derivative events (next section) in generating human isoform sequence diversity. However, the mechanistic basis of how these features will correspond to traditional AS events is not known, nor the nature of Block-I differences, which are more apparent in lower sID bins and could be caused by exon locus undergoing amino acid change or transitioning from UTR/CDS while maintaining their genomic coordinates.

### Analyzing AS events effect on mIDI isoform divergence

The previous analysis divulged role of block-I and block-III attributes and highlight their contribution along with compositionally differing exon loci in generating isoform diversity. However, it does not provide the specific type(s) of AS event or their combination(s) association with isoform divergence. To investigate this, we extended the analyses to examine profile of AS events, preponderant in leading to mIDI construct from RISO. ENACT enables this analysis through exon Block segregation of information and their locus based tracking across isoforms facilitating aim to relate AS event(s) with distinction of CDS/UTR regions (see methods). Utilizing this, we identified four well established primary AS events, (Exon gain (EG) and Exon skipping (ES), Alternate splice site change (A(ss)) and Intron retention (IR)). We pinpointed subtypes of ES/EG provided they alter at initial and terminal exon index of GA, as Alternate first exon (AFE) and Alternate last exon (ALE) events. For exons undergoing alternate sequence adaptations, we defined Sequence variation 1 and 2 (Sv1 and Sv2 events), depicting gain of residues and frame change events, respectively.

Initially, we assessed the prevalence of single AS events (Figure 4A) occurring in both coding/non-coding exons in mIDI comparison to RISO for all human genes (GeneISF2), followed by a preponderance of two combined AS events. We computed gene fractions undergoing at least one of such AS events (Figure 4) and among them ES (Exon Skipping) (Figure 4A) is predominant in coding regime, followed by Alternate splice site (A(ss)). AFE is commonly observed in the non-coding regime (Figure 4B). Investigating their prevalence previously considered sID bins (Figure 4A1-5), we observed that, for highly divergent isoforms (sID < 50%), ALE, Sv2, and IR events were predominant and vary the most from the expected distribution (see supplementary section S2). ES events were predominant in 50-90% sID bins, while A(ss) event is dominant in 90-100% sID. Among non-coding exons, AFE is more common in 100% sID. Detailing this further for two combined AS events (Figure 5) across sID bins showed that for coding regime, AFE-ALE, and ’AFE-Sv2’ were frequent in low sID (0-50%). We also observe that A(ss)-ES, and ALE-ES, AFE-ES are prevalent in high sID (50-100%). This suggests exon skipping or alternate splice site frequently combined with alternate translational start/end to bring isoform diversity.

**Figure 4:**
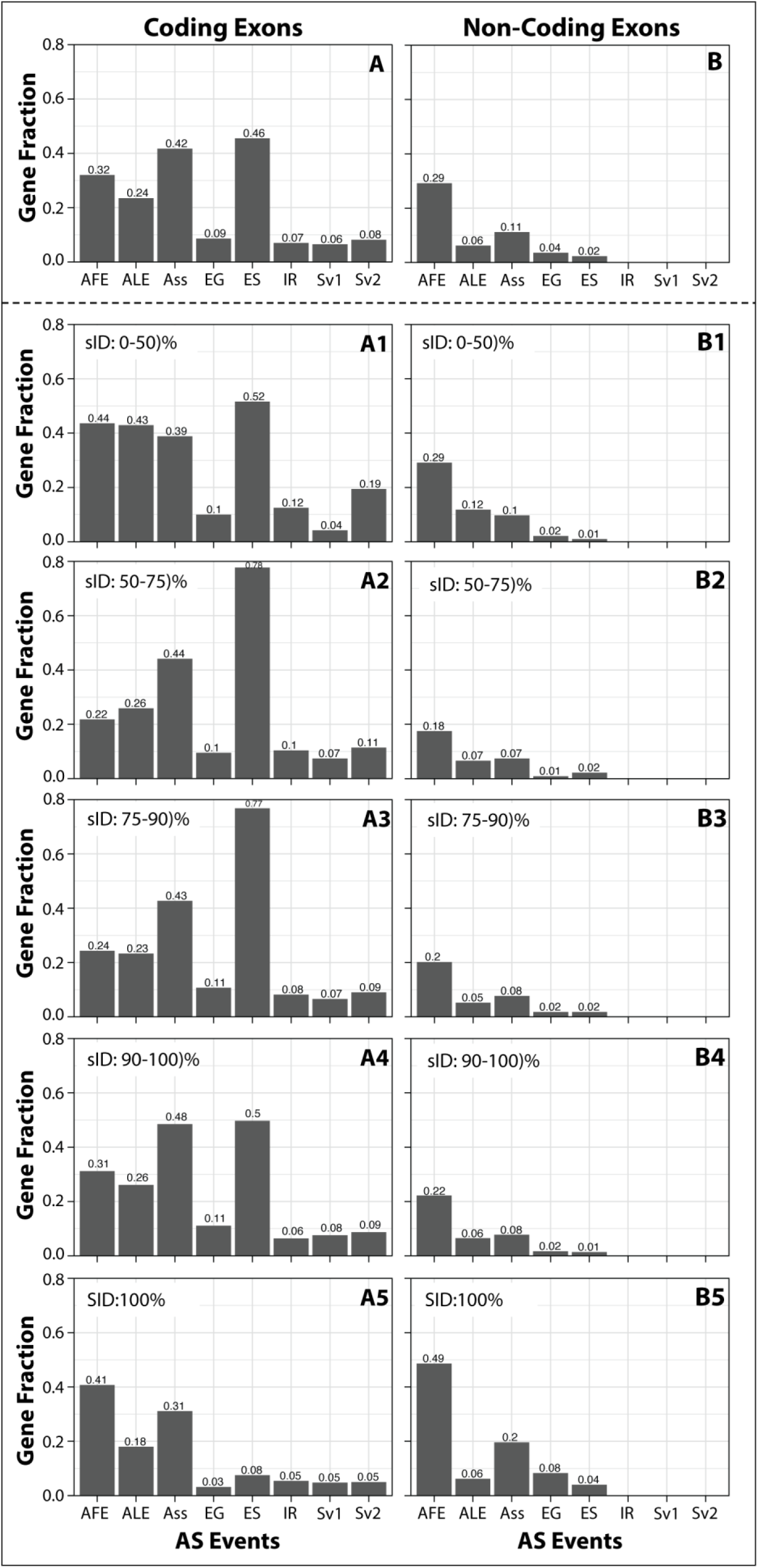
Occurrence of single AS event in mIDI/RISO for coding and non-coding exons. The histogram showing the gene fraction observed for at least one AS event observed in mIDI involving either coding exons (left column, A) or non-coding exons (right column, B). The top rows in A and B panels are for the occurrence of AS events computed for the human genes dataset. The vertical panels from A1-A5 or B1-B5 show the gene fraction observed for an AS events in various sID bins (0-50%), 50-75%), 75-90%), 90-100%), and 100%. The gene fraction of events in each cell has been normalized to the total genes in that cell (bin), and an AS event of a gene is counted only once. The following AS events are considered: Alternate First Exon (AFE), Alternate Last Exon (ALE), Alternate splice site (A(ss)) choice, exon gain (EG) in mIDI, exon skipping (ES), intron retention (IR) and, Sv1 and Sv2 have identical Blocks-II and III features but vary in Block-I by no amino acid sequence change and frame-shift change respectively.

**Figure 5:**
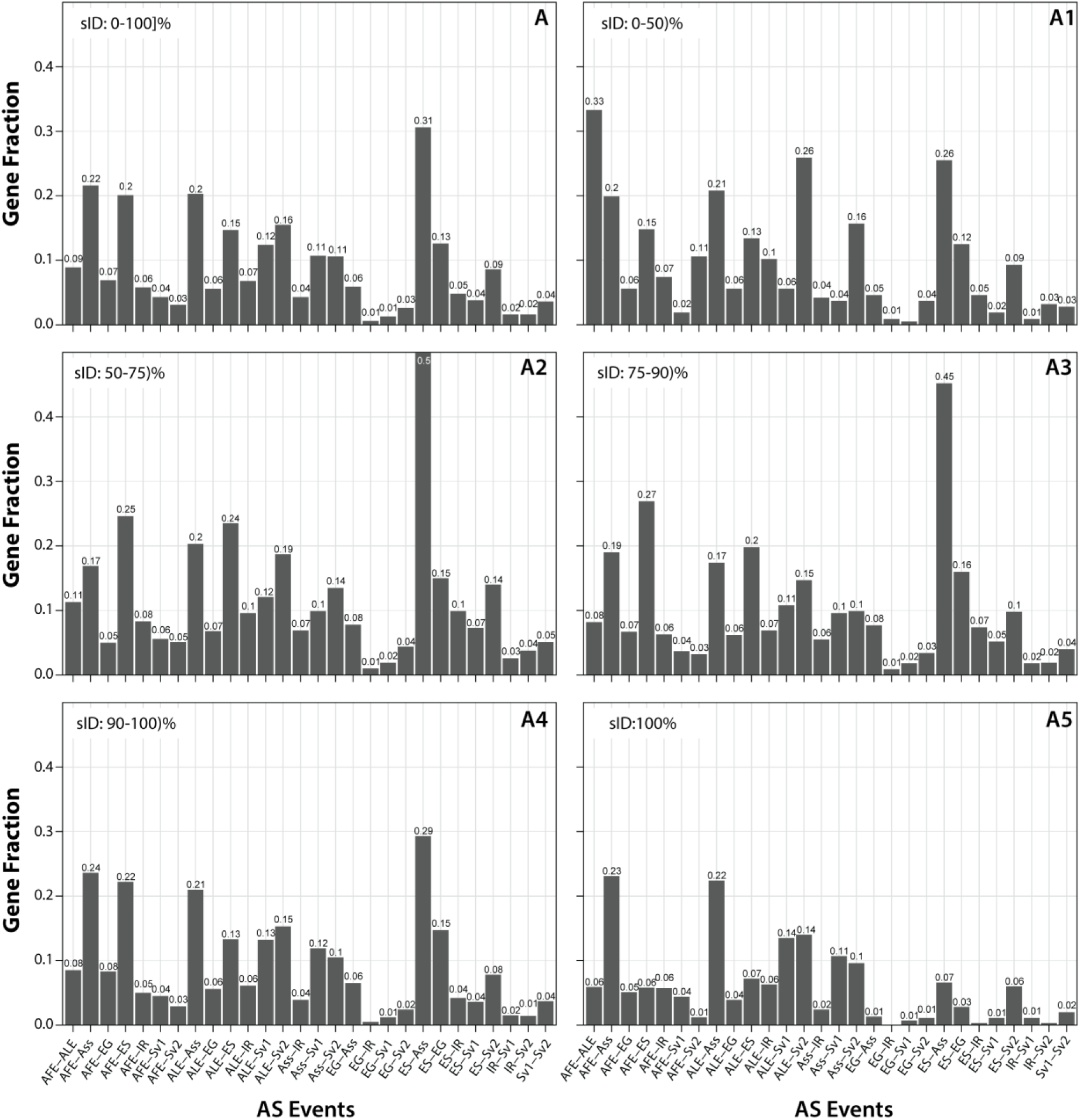
Occurrence of two AS combined events in mIDI/RISO for coding and non-coding exons. The histograms show the distribution of gene fractions observed for combined events involving at least two distinct AS subtypes in mIDI/RISO. Panel A shows the distribution of two AS event occurrences computed for the whole dataset. The plots A1-A5 is distribution of combined events in various sID bins (0-50%), 50-75%), 75-90%), 90-100%), and 100%. The gene fraction of events in each cell has been normalized to the total genes in that cell (bin), and a combined AS event of a gene is counted only once. Total genes used per cell as denominator, pre-filters to involve only those genes, which are undergoing atleast 1 of above event combination with 1-member event affecting genic coding region. The following AS events are considered: Alternate First Exon (AFE), Alternate Last Exon (ALE), Alternate splice site (A(ss)) choice, exon gain (EG) in mIDI, exon skipping (ES), intron retention (IR) and, Sv1 and Sv2 have identical Blocks-II and III features but vary in Block-I by no amino acid sequence change and frame-shift change respectively.

In summary, analysis of exon composition and similarity between mIDI and RISO, highlighted broadened roles of exonic splice site (Block-III) and sequence variations (Block-I) in influencing the spectrum of isoform diversity. Examining these observations in context of specific AS events, we noted predominance of alternate translational start/end (AFE/ALE), A(ss) and ES subtypes. Considering these insights and documented higher incidences of exon skipping, introspection of affected localized region by exons undergoing such events seems paramount. Moreover, co-role of A(ss) events and their association with ES are also largely unknown. Taking into account frequent association of A(ss) also with AFE / ALE, we hypothesized that they might be preponderant towards the start and end of coding gene architecture.

### Location of ES/A(ss) event in gene architectural context

Previous analyses highlighted broad roles of ES/A(ss)/AFE/ALE in generating isoform diversity. To explore the potential association between exonic loci undergoing these events and their linear spatial indexing within gene architecture, we conducted an analysis to ascertain their prevalence across coding and noncoding regimes of gene architecture and their intersecting regions. ENACT attributes of concerned loci were related to linear index and we calculated spatial positioning by normalizing ‘aa’ length contributed till exonic loci of interest with total ‘aa’ between first and last coding exons. Quantified spatial indexing correspondingly has been binned to intervals of 20% and relative proportion of exonic loci has been illustrated for ES and A(ss) events in Figure 6. It shows that ES event undergoing exonic loci are predominantly present in the 5’UTR region. Conversely, prevalence of such loci to undergo A(ss) event are comparatively scarce. Interestingly, occurrence of loci’s undergoing ES and A(ss) exons are relatively abundant in the first 20% and last 20% of gene architecture (Figure 6). This observation suggests that protein termini tend to be relatively more susceptible to inclusion/exclusion of exons or their n/c/b modifications, with possible implications to change protein sequence. Predominance of ES events in animals has been noted (36), however not their association to co-occur frequently with A(ss) events. Above observations of these events to spatially localize to start and end of coding gene architecture, affirms their potential role in facilitating alternate translation initiation and termination sites while purposing ES and A(ss) event definitions. This also highlights crucial amalgamated role of splicing complex events extending to other post transcriptional processes and showcase their synergistic contribution in generating proteogenomic diversity.

**Figure 6:**
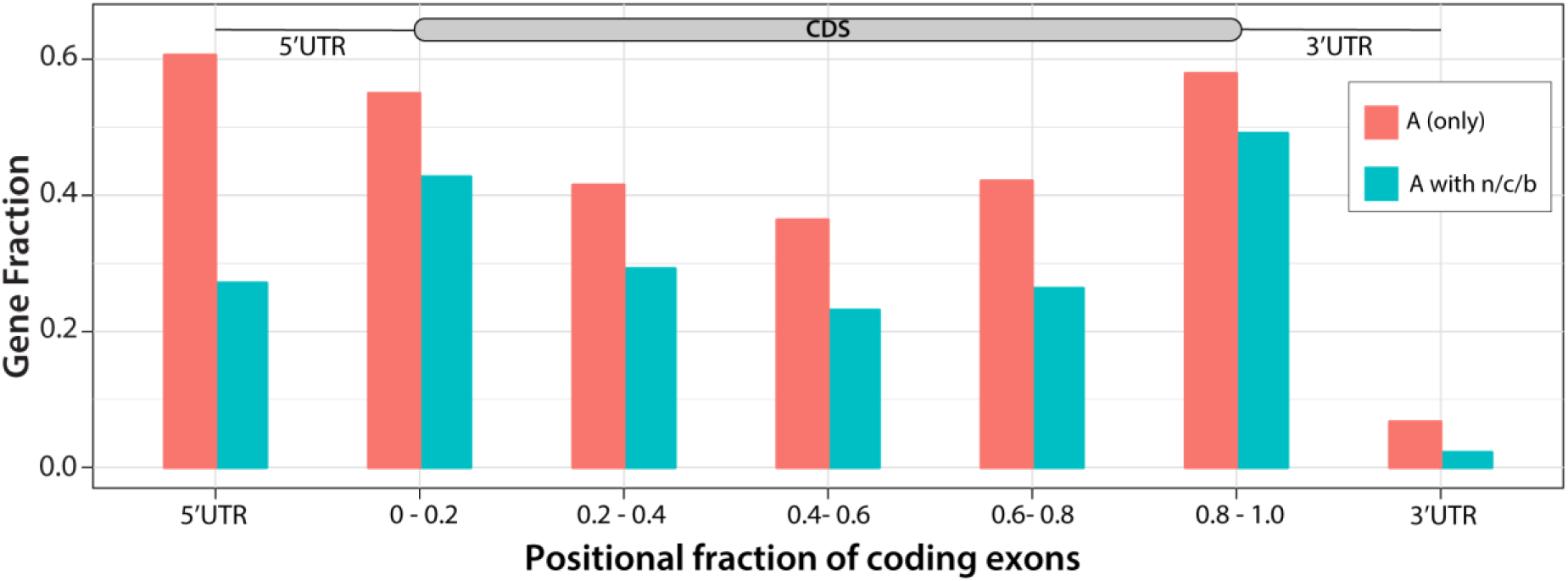
Occurrence and prevalence of alternate exons (with and without splice site variation) in genic architectural position. The histogram showing distribution of gene preponderance of having exons undergoing A(ss) (without n/c/b) and A(ss) (with n/c/b) instances. We identified genes exhibiting at least one coding exonic loci undergoing alternate splice site (n/c/b) choice instances. Positional indexing of exonic loci (alternate loci undergoing or not undergoing A(ss)) were calculated (see text) and categorized into five bins of 0.2 width and 5’/3’ UTR regions (outside first and last coding exon). Gene fraction (Y axis) depicts relative gene proportion from considerable set harboring atleast 1 exonic loci in specified bin range undergoing particular event. 10389 genes (considerable gene set for ’A with n/c/b’ event type’), have at least one exon with T/D tag in Block-I and n/c/b in Block-III (F-tag in Block-II is considered as A(ss)), out of which 9264 genes (considerable gene set for ‘A (only)’ event type) comprised Block-III without n/c/b but Block-II under alternate attribute.

### Functional diversity in different identity bins illustrated by ENACT nomenclature

To demonstrate the functionally induced variation as function of the previously discussed sID and sCOV analysis, we selected four representative examples having isoforms diverging in sID from RISO. We also illustrate the utility of exon-centric annotation in understanding the functional diversity introduced in those from RISO.

#### A. ADAM8

The ADAM8 gene encodes a protein belonging to the membrane-anchored disintegrin and metalloprotease proteinases family that cleaves the extracellular domain of several cell surface proteins and receptors (37). The ADAM8 protein is involved in various cellular functions such as inflammation, immunomodulation, neutrophil activation/mobility, immune cell migration, osteoclast stimulating factor, and neurodegeneration (38–40). The ADAM8 domain architecture consists of an N-terminal prodomain, a catalytic metalloproteinase domain, a disintegrin domain involved in interaction with integrins, a cysteine-rich domain followed by a transmembrane region, and a C-terminal domain probably involved in protein-protein interaction through SH3 or proline-rich regions (41).

The ADAM8 gene comprises 23 coding exons, one each of non-coding and dual exons. Of the coding exons, 17 are constitutive/constitutive-like, and the rest are alternate. We analyzed three well-annotated isoforms harboring a combination of AS events (Figure 7A). The reference isoform (IS-1; NP_0011003.3) has Pep_M12B_propep, Reprolysin (metalloproteinase), Disintegrin, and ADAM_CR (cysteine-rich domain) Pfam domains lying before transmembrane region (seen in ENACTdb). The skipping of exon-21 is combined with 5ss of exon-22 (ES/A(ss) linear in sequence index) (IS-2: NP_001157961.1), leading to frame shift in subsequent exons and premature termination in exon-24. These could be noted by their EUIDs: T.2.F.22.n.1, T.2.G.23.0.0, and T.2.A.24.0.0. Thus, resulting isoform lacks the proline-rich region required for protein-protein interaction. The IS-2 isoform is expressed in metastatic lung cancer cell lines (41). The IS-3 (mIDI, NP_001557962.1, sID: 82.2%, sCOV:88.1%) exhibit skipping of exons 2 to 4, which affects the local reading frame involving exons 5 to 7. However, the reading frame is restored from exon 8 onwards by 5ss AS event (Figure 7A). Due to a change in the amino acid sequence in IS-3, the pro-domain cannot be identified in the N-terminal region of this isoform, suggesting that it may have constitutive metalloproteinase activity. Furthermore, IS-3 also lacks 2 out of 4 glycosylation sites, which are part of the pro-domain (42). However, one of the conserved Glutamate (158E) essential for the catalytic removal of the pro-domain (43) is preserved in IS-3. Further experimental studies would provide information on the enzymatic activity and biological role of IS-3.

**Figure 7:**
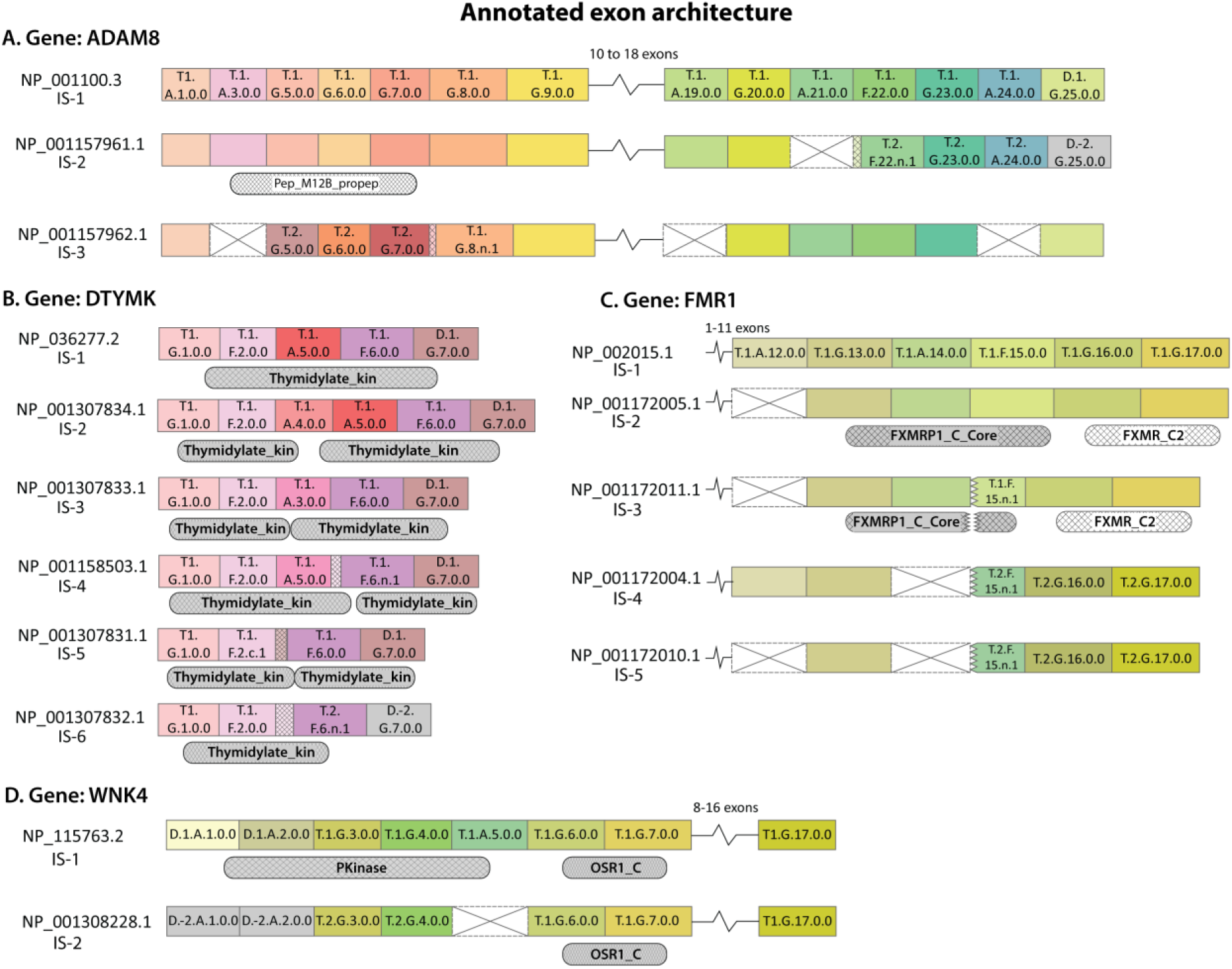
ENACT annotation of ADAM8, DTYMK, FMR1 and WNK4 isoforms and exons. The NCBI identifier is shown for each isoform, and exons are represented as colored rectangle boxes with their respective EUID. The absence or skipped exon is shown with a crossed empty rectangle box. The small extension of exon rectangle boxes with crisscross filled lines represents extended exon boundaries due to alternate splice sites (5ss/3ss). If an exon does not show variation with exon(s) in the isoform shown above, then the EUID is not labelled. The break shows that exons intervening in the region do not change in the isoforms. The jagged edges of a rectangle represent 5’ss or 3’ss alternate splice sites. The isoforms sharing Pfam domains for a region are shown below exon layout. In ADAM8 isoforms, IS-3 lacks the Pep_M12B_propep Pfam domain. All isoforms of DTYMK consist of shortened or extended. Thymidylate kinase domain. In the FMR1 gene, FXMRP1_C_Core and FXMR_C2 domains are assigned in the same region of IS-1 and IS-2. However, IS-4 and IS-5 lack these two domains. Similarly, IS-2 of WNK4 lacks Pkinase domain in the N-terminal region.

#### B. DTYMK

The deoxythymidylate kinase (DTYMK) gene encodes an enzyme essential for DNA synthesis, nuclear genome stability, and mitochondrial copy number maintenance (44). The enzyme catalyzes the transfer of γ-phosphate of ATP to dTMP in the presence of Mg^2+^ ion. The gene expression peaks during the S-phase and is low from mitosis to the early G-1 phase (44). The gene is upregulated in cancer cells (45) and tissues linked to the development of neurodegenerative diseases similar to severe microcephaly in humans (46). Previous studies on DTYMK sequences have identified three sequence motifs: a) lid region (residue 142 - 154) required for conformation change and P loop motif; b) the DRX motif (X = Y/F, D) involved in catalysis and c) non-covalent interaction network formed by R76, D96 and π-π stacking of residues F72, F105 and Y151 (44). These residue numbers correspond to the protein sequence of NCBI protein id NP_036277.2. The DTYMK gene consists of six coding and one dual exon. Exons 3, 4, and 5 of six coding are alternate exons. The Pfam thymidylate kinase domain could be associated with all six isoforms (Figure 7B). However, it either was an extended domain in some isoforms or atrophied in others. For instance, exon 4 (EID: T.1.A.4.0.0) occurs only in IS-2 (NP_001307834.1) resulting in an insertion region within the Thymidylate kinase domain.

The shortest isoform (IS-6) is 113 amino acids long, primarily due to ES events involving exons 3 to 5 with amino acid sequence variation in exon-6 due to 5ss (EUID: T.2.F.6.n.1) and premature termination. Interestingly, the exon alignment view shows that exons 3 and 4 are mutually exclusive AS events, as both do not co-occur in any isoform. Since alternate exons are present in the middle of the DTYMK gene, we examined the conservation of structural and sequence features among isoforms. In recent work, Frisk et al. (44) investigated changes in the domains of thymidylate kinase and found that isoforms lacking exons 3, 4, 5, or 6 with sequence variation lack crucial sequence motifs required for DTMYK activity. The reference isoform IS-2 (NP_001307834.1) has the insertion of 39 residues and weak enzymatic activity. Notably, IS-2 is expressed along with IS-1 (NP_036277.2) in fibroblast cell lines, suggesting that IS-2 may affect IS-1 function by sequestering substrate for enzyme activity, as it has conserved binding site residues. IS-5 (mIDI, NP_001307831.1, sID:67.3%, sCOV:67.3%) has undergone ES of 4 and 5 exons but after alternate 3’ splice site choice for exon 2, leading to sID and sCOV of only 67.3%, likely from altered integrity of second signature of Pfam thymidylate kinase domains. However, this and other isoforms and their complex interactions, structural fate, and functional divergence have not yet been elucidated experimentally.

#### C. Fragile X messenger ribonucleoprotein 1 (FMR1)

The FMR1 gene encodes fragile X mental retardation protein (FMRP), whose loss of function causes inheritable disorder fragile X syndrome and premature ovarian failure (47). The protein regulates the translation of a subset of mRNAs and the shuttling of mRNAs in the intracellular compartment. As a result, it is usually localized in polyribosomes or ribonucleoprotein complexes. Although FMR1 is expressed in almost all tissues, it is abundant in testes and brain cells. In neuronal cells, it regulates synaptic plasticity (48). The gene has 17 coding exons, which undergo alternative splicing to produce multiple transcripts that could vary up to 49 isoforms (49). Some human and mouse isoforms are well characterized by their sequence features and role in function or cellular localization (50,51).

We analyze representative ENACT annotated isoforms of FMR1, focusing on AS events at the C-terminus to explore possible effects of alternative splicing on protein function. The FMRP protein is an RNA-binding protein consisting of RNA-binding domain, i.e., hnRNP K Homology (KH) domains and RGG box motif. Two KH domains span exons 3-8, and the RGG motif is located in exon 15. The reference isoform IS-1 (NP_002015.1) consists of all 17 coding exons and two C-terminal Pfam domains, FXMRP1_C_Core and FXMR_C2 span exons 13-15) and exons 16-17, respectively (Figure 7C). Exon 12 (T.1.A.12.0.0) is skipped without affecting the domain architecture in IS-2 (NP_001172005.1) with no consequence on the features of the protein domain. On the contrary, exon 15 undergoes identical 5ss in IS-3, IS-4, and IS-5 and ES events simultaneously in those isoforms but with varying consequences on the amino acid sequence of exon 15, as evident from subsequent EUIDs in these isoforms. In mIDI (IS-3: NP_001172011.1, sID:92.7%, sCOV:92.7%), the 5ss AS event of exon 15 is assigned with EUID: T.1.F.15.n.1, showing that its N-terminal sequence is altered, which results in truncation of FXMRP1_C_Core domain. However, in IS-4 (NP_001172004.1, sID:100% and sCOV:67%) and IS-5 (NP_001172010.1, sID:95% and sCOV:65%), the exon 14 skipping event leads to the reading frame shift, resulting in the amino acid sequence change from exons 15-17. The amino acid change is evident in their nomenclature, as exons 15, 16, and 17 are assigned EUIDs T.2.F.15.n.1, T.2.G.16.0.0, and T.2.G.17.0.0 respectively. The second character (T.2.) shows that the amino acid sequence is different from the reference exon without any change in their genomic coordinates. Due to the frameshift of the reading frame and loss of exon 14 in IS-4 and IS5, the two Pfam domains are lost and may lead to altered protein function. In addition, exon 14 harbors a nuclear export signal, and in the absence of this, IS-4 and IS-5 will be unable to perform nucleocytoplasmic shuttling functions.

#### D. WNK4 gene

The WNK4 gene belongs to the conserved ‘With no lysine (WNK)’ group of serine/threonine kinases (STK) in eukaryotic organisms. These have been named because of their atypical positioning of catalytic lysine in subdomain II instead of I, as in other STKs. WNK4 is expressed primarily in the kidney. This with other members of the family has role in modulating the balance between sodium chloride re-absorption and renal potassium ion secretion (52) by regulating the activities of cation-coupled cotransporters (SLC12, NCC), ion channels (ENaC) and ion exchangers (53,54). The mutation in the WNK4 gene is associated with a rare genetic type of hypertension called pseudohypoaldosteronism type 2 (PHA2). The protein has two Pfam domains (Protein kinase and Oxidative-stress-responsive Kinase1 C-terminal domain), while the rest of the sequence is intrinsically disordered. The OSR1_C encompasses Pask-Fray 2 (PF2) domain, which is known to interact with RFX[VI] motif and suppress the activity of kinase domain (52).

A total of 13 isoforms are listed in the NCBI RefSeq database. Of these, two isoforms are reviewed, *i.e.,* having experimental validation (Figure 7D). Among the 19 exons, the first two are ‘Dual’ as they are coding in some transcripts and non-coding in others. These two exons are encoded as coding in the reference isoform (NP_115763.2), resulting in Pfam Protein kinase domain (Figure 7D) being assigned to the N-terminal region of the protein. In contrast, the first two exons are non-coding in IS-2 (NP_001308228.1, sID: 94.9% sCOV:68.5% ) with alternate translation initiation site in exon 3 (EUID: T.2.G.3.0.0), leading to a change in amino acid of exons 3-4 due to frameshift (EUIDs are T.2.G.3.0.0 and T.2.G.4.0.0, evident from second character). Interestingly, the exon 5 skipping (ES) AS event restores the reading frame and the rest of protein sequence is maintained as in the reference isoform, which is also evident from the EUIDs of exons. IS-2 shows kinase domain loss, but supports OSR1_C (PF2) domain. The latter domain is suggested to interact with SPAK/OSR1 protein, suggesting that IS-2 may act as a sequestering factor for them and affect their biological function.

## Discussion

In the present work, we have systematically investigated the roles of various splicing events and their combination leading to isoform diversity. RNAseq studies have hallmarked the role of splicing introduced changes in transcriptome diversification. Our work here extends impact of splicing and related processes by studying their impact on imparting intra gene protein sequence diversity.

In our study, we have formulated an innovative framework ENACT, to annotate exons with unique attributes, where relative position and splice site variants were extracted from GC and translational potential and variability from CGC. These exon attributes were depicted using a single character and grouped in three Blocks. Employing this framework, we investigated the isoform diversity of 14761 human genes having at least two protein-coding isoforms. We identified and compared attributes differing between mIDI and RISO for different sID bins, to observe differing contribution of exon translational features (Block-I) and splice site choices (Block-III) in them. Impact of Block-I and Block-III attribute variability were comparable in low sequence divergence (high sID) but showed contrast with bias towards Block-I attributes in high sequence divergence (low sID). When we further deduced AS events as a function of exon feature Block combination, distinct preponderance of events were noted in spectrum of sID bins, where AFE was favored in UTR region for higher sID intervals and the combination of AFE/ALE were favored in CDS for lower sID intervals. Combination of ES/A(ss) events among coding region were also noted for similar distribution in most sID bins and their subsequent spatial indexing on gene architecture unveiled them to be localized often to UTR regime and termini of coding gene architecture. Concomitantly, how their combination might be impacting functional scope of resulting isoform in addition to influence alternate translation start and termination were illustrated through four representative examples. Although the current study is limited to curated gene models, however it serves well to concur with aspects of alternative splicing observed in literature that is discussed briefly:

a. Studies have determined several complexes AS event types / higher entropy events and associated them with the states of pathologies / specialized processes (27,28). However, most of them were present near the 5’UTR or affected the alternate splice sites of exonic loci. Our work finds events associated with imparting variation due to ATLI/ATLT.
b. The scope of an alternate fraction of nucleotides being largely contributed by alternate transcription has been suggested previously (20). However, lacked any follow-up studies to determine event types. Our study affirms the ES events being prevalent at termini of coding gene architecture, but also observes the enrichment of many loci at those positions undergoing splice site choices. In separate analysis (data not shown, manuscript under preparation), we observed an emerging gene population preferring that exonic locus to introduce alternate translational features.
c. Recent literature has also emphasized exon mediated activation of transcription start and end (7) and existence of hybrid internal terminal exons (29) with the potential to alter the reading frame. Our study finds curated transcript models and ENACT framework implied transcript annotations (case studies), that clearly show potential realizations of such exons to instill protein diversity. They emphasize how combinations of AS events, such as (A(ss) and ES) distinctly in distant positions of transcript structure, alters/introduce functional changes reminiscent of diversity from reference isoform. Additionally, case studies also reflect on understudied synergistic impact of such event combinations to rescue altered transcript (by skipping of non 3n exon, or alternate uORF preference) by restoring its partial composition to that of RISO.

Considering the phenotypic complexity of higher eukaryotes, the role of alternative splicing in influencing transcriptomic/translational processes of genes to generate isoform diversity needs detailed investigation. Although extent of RNA-seq analysis addresses answers many of transcriptome diversity concerns, relevance on their translated counterparts are only rudimentary understood. The ability to integrate transcriptome and proteome information would be helpful to understand complete landscape of AS. The ENACT framework helps to provide a nuanced understanding of gene and transcript-specific regional translational potential while tracking exons. Further, the protein-specific information, such as secondary structure, and sequence domains, can assist in analyzing the effect of exon inclusion/exclusion and their splice site variations on protein functions. We have restricted the present work to five well-studied model organisms. Future work on other organisms will clarify the extent of regional sub-components (Blocks of ENACT entities) and their evolutionary dynamics, with the scope to functionally delineate the alternate transcription/ translational/ splicing processes.

## Supporting information

SuppDataExcel

## Data Availability

All data generated during this study are publicly available at: www.iscbglab.in/enactdb/. Associated algorithmic procedures have been described in methods.

## Supplementary data

PDF file enclosing supplementary data attached.

## Author contribution

Conceptualization: SBP Methodology: PV, DT, SBP

Investigation: PV, DT

Visualization: PV, DT, SBP

Supervision: SBP

Writing—original draft: PV, DT

Writing—review & editing: PV, DT, SBP

## Acknowledgement

The authors acknowledge Deepanshi Awasthi for her critical reading of the manuscript. We also acknowledge computing facility Param Smriti formed under National Supercomputing Mission.

## Funding

IISER Mohali

Bioinformatics Center (BT/PR40419/BTIS/137/36/2022), Department of Biotechnology under the Ministry of Science and Technology, Govt. of India

National Network Project (BT/PR40198/BTIS/137/56/2023), Department of Biotechnology under the Ministry of Science and Technology, Govt. of India

### Box-1 ENACT Algorithm

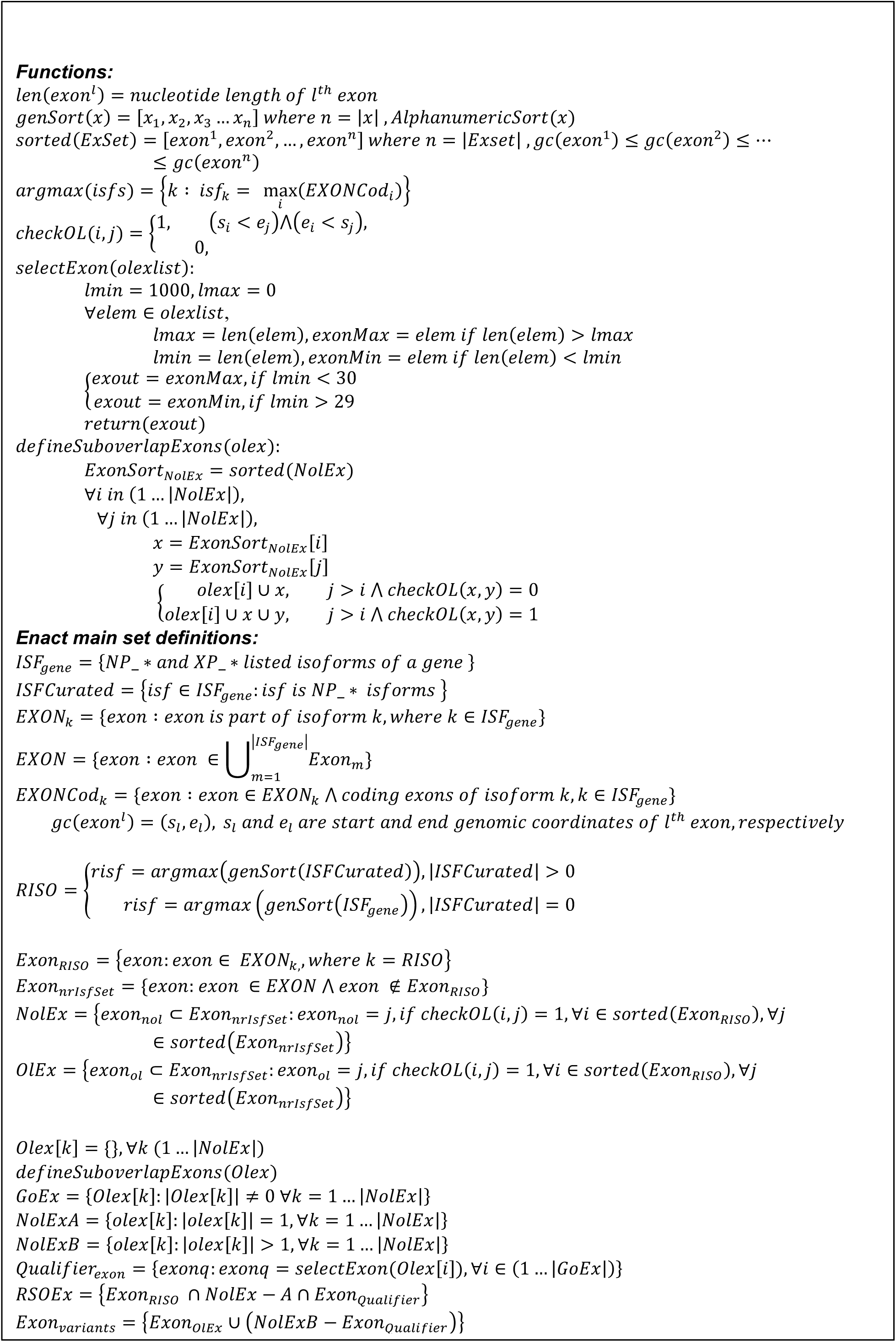

## References

1. Verta, J.P. and Jacobs, A. (2022) The role of alternative splicing in adaptation and evolution. Trends Ecol Evol, 37, 299–308.

2. Tapial, J., Ha, K.C.H., Sterne-Weiler, T., Gohr, A., Braunschweig, U., Hermoso-Pulido, A., Quesnel-Vallières, M., Permanyer, J., Sodaei, R., Marquez, Y. et al. (2017) An atlas of alternative splicing profiles and functional associations reveals new regulatory programs and genes that simultaneously express multiple major isoforms. Genome Research, 27, 1759–1768.

3. Baralle, F.E. and Giudice, J. (2017) Alternative splicing as a regulator of development and tissue identity. Nature reviews Molecular cell biology, 18, 437–451.

4. Rogalska, M.E., Vivori, C. and Valcárcel, J. (2023) Regulation of pre-mRNA splicing: roles in physiology and disease, and therapeutic prospects. Nat Rev Genet, 24, 251–269.

5. Ni, T., Corcoran, D.L., Rach, E.A., Song, S., Spana, E.P., Gao, Y., Ohler, U. and Zhu, J. (2010) A paired-end sequencing strategy to map the complex landscape of transcription initiation. Nat Methods, 7, 521–527.

6. Kamieniarz-Gdula, K. and Proudfoot, N.J. (2019) Transcriptional Control by Premature Termination: A Forgotten Mechanism. Trends Genet, 35, 553–564.

7. Fiszbein, A., Krick, K.S., Begg, B.E. and Burge, C.B. (2019) Exon-Mediated Activation of Transcription Starts. Cell, 179, 1551–1565 e1517.

8. Bentley, D.L. (2014) Coupling mRNA processing with transcription in time and space. Nat Rev Genet, 15, 163–175.

9. Kornblihtt, A.R., Schor, I.E., Allo, M., Dujardin, G., Petrillo, E. and Munoz, M.J. (2013) Alternative splicing: a pivotal step between eukaryotic transcription and translation. Nat Rev Mol Cell Biol, 14, 153–165.

10. Braunschweig, U., Gueroussov, S., Plocik, A.M., Graveley, B.R. and Blencowe, B.J. (2013) Dynamic integration of splicing within gene regulatory pathways. Cell, 152, 1252–1269.

11. Shabalina, S.A., Spiridonov, A.N., Spiridonov, N.A. and Koonin, E.V. (2010) Connections between alternative transcription and alternative splicing in mammals. Genome Biol Evol, 2, 791–799.

12. Kochetov, A.V. (2008) Alternative translation start sites and hidden coding potential of eukaryotic mRNAs. Bioessays, 30, 683–691.

13. Johnstone, T.G., Bazzini, A.A. and Giraldez, A.J. (2016) Upstream ORFs are prevalent translational repressors in vertebrates. EMBO J, 35, 706–723.

14. Lee, S., Liu, B., Lee, S., Huang, S.X., Shen, B. and Qian, S.B. (2012) Global mapping of translation initiation sites in mammalian cells at single-nucleotide resolution. Proc Natl Acad Sci U S A, 109, E2424–2432.

15. Ingolia, N.T., Brar, G.A., Stern-Ginossar, N., Harris, M.S., Talhouarne, G.J., Jackson, S.E., Wills, M.R. and Weissman, J.S. (2014) Ribosome profiling reveals pervasive translation outside of annotated protein-coding genes. Cell Rep, 8, 1365–1379.

16. Ji, Z., Song, R., Regev, A. and Struhl, K. (2015) Many lncRNAs, 5’UTRs, and pseudogenes are translated and some are likely to express functional proteins. Elife, 4, e08890.

17. Sudmant, P.H., Lee, H., Dominguez, D., Heiman, M. and Burge, C.B. (2018) Widespread Accumulation of Ribosome-Associated Isolated 3’ UTRs in Neuronal Cell Populations of the Aging Brain. Cell Rep, 25, 2447–2456 e2444.

18. Kesner, J.S., Chen, Z., Shi, P., Aparicio, A.O., Murphy, M.R., Guo, Y., Trehan, A., Lipponen, J.E., Recinos, Y., Myeku, N. et al. (2023) Noncoding translation mitigation. Nature, 617, 395–402.

19. Aspden, J.L., Wallace, E.W.J. and Whiffin, N. (2023) Not all exons are protein coding: Addressing a common misconception. Cell Genom, 3, 100296.

20. Shabalina, S.A., Ogurtsov, A.Y., Spiridonov, N.A. and Koonin, E.V. (2014) Evolution at protein ends: major contribution of alternative transcription initiation and termination to the transcriptome and proteome diversity in mammals. Nucleic Acids Res, 42, 7132–7144.

21. Deveson, I.W., Brunck, M.E., Blackburn, J., Tseng, E., Hon, T., Clark, T.A., Clark, M.B., Crawford, J., Dinger, M.E., Nielsen, L.K. et al. (2018) Universal Alternative Splicing of Noncoding Exons. Cell Syst, 6, 245–255 e245.

22. Baek, D., Davis, C., Ewing, B., Gordon, D. and Green, P. (2007) Characterization and predictive discovery of evolutionarily conserved mammalian alternative promoters. Genome Res, 17, 145–155.

23. Derti, A., Garrett-Engele, P., Macisaac, K.D., Stevens, R.C., Sriram, S., Chen, R., Rohl, C.A., Johnson, J.M. and Babak, T. (2012) A quantitative atlas of polyadenylation in five mammals. Genome Res, 22, 1173–1183.

24. Foissac, S. and Sammeth, M. (2007) ASTALAVISTA: dynamic and flexible analysis of alternative splicing events in custom gene datasets. Nucleic Acids Research, 35, W297–W299.

25. Sammeth, M., Foissac, S. and Guigó, R. (2008) A general definition and nomenclature for alternative splicing events. PLoS Comput. Biol., 4, e1000147.

26. Xing, Y., Yu, T., Wu, Y.N., Roy, M., Kim, J. and Lee, C. (2006) An expectation-maximization algorithm for probabilistic reconstructions of full-length isoforms from splice graphs. Nucleic Acids Res, 34, 3150–3160.

27. Vaquero-Garcia, J., Barrera, A., Gazzara, M.R., Gonzalez-Vallinas, J., Lahens, N.F., Hogenesch, J.B., Lynch, K.W. and Barash, Y. (2016) A new view of transcriptome complexity and regulation through the lens of local splicing variations. Elife, 5, e11752.

28. Sterne-Weiler, T., Weatheritt, R.J., Best, A.J., Ha, K.C.H. and Blencowe, B.J. (2018) Efficient and Accurate Quantitative Profiling of Alternative Splicing Patterns of Any Complexity on a Laptop. Mol Cell, 72, 187–200 e186.

29. Fiszbein, A., McGurk, M., Calvo-Roitberg, E., Kim, G., Burge, C.B. and Pai, A.A. (2022) Widespread occurrence of hybrid internal-terminal exons in human transcriptomes. Sci Adv, 8, eabk1752.

30. Reixachs-Solé, M. and Eyras, E. (2022) Uncovering the impacts of alternative splicing on the proteome with current omics techniques. Wiley Interdiscip. Rev. RNA, 13, e1707.

31. Tsang, M.J. and Cheeseman, I.M. (2023) Alternative CDC20 translational isoforms tune mitotic arrest duration. Nature, 617, 154–161.

32. O’Leary, N.A., Wright, M.W., Brister, J.R., Ciufo, S., Haddad, D., McVeigh, R., Rajput, B., Robbertse, B., Smith-White, B., Ako-Adjei, D. et al. (2016) Reference sequence (RefSeq) database at NCBI: current status, taxonomic expansion, and functional annotation. Nucleic Acids Res, 44, D733–745.

33. De Conti, L., Baralle, M. and Buratti, E. (2013) Exon and intron definition in pre-mRNA splicing. Wiley Interdiscip Rev RNA, 4, 49–60.

34. O’Leary, N.A., Wright, M.W., Brister, J.R., Ciufo, S., Haddad, D., McVeigh, R., Rajput, B., Robbertse, B., Smith-White, B., Ako-Adjei, D. et al. (2016) Reference sequence (RefSeq) database at NCBI: current status, taxonomic expansion, and functional annotation. Nucleic Acids Res., 44, D733–745.

35. Cock, P.J., Antao, T., Chang, J.T., Chapman, B.A., Cox, C.J., Dalke, A., Friedberg, I., Hamelryck, T., Kauff, F. and Wilczynski, B.J.B. (2009) Biopython: freely available Python tools for computational molecular biology and bioinformatics. 25, 1422.

36. Shabalina, S.A., Ogurtsov, A.Y., Spiridonov, N.A. and Koonin, E.V. (2014) Evolution at protein ends: major contribution of alternative transcription initiation and termination to the transcriptome and proteome diversity in mammals. Nucleic Acids Res., 42, 7132–7144.

37. Fourie, A.M., Coles, F., Moreno, V. and Karlsson, L. (2003) Catalytic activity of ADAM8, ADAM15, and MDC-L (ADAM28) on synthetic peptide substrates and in ectodomain cleavage of CD23. J. Biol. Chem., 278, 30469–30477.

38. Yamamoto, S., Higuchi, Y., Yoshiyama, K., Shimizu, E., Kataoka, M., Hijiya, N. and Matsuura, K. (1999) ADAM family proteins in the immune system. Immunology Today, 20, 278–284.

39. Schlomann, U., Rathke-Hartlieb, S., Yamamoto, S., Jockusch, H. and Bartsch, J.W. (2000) Tumor necrosis factor alpha induces a metalloprotease-disintegrin, ADAM8 (CD 156): implications for neuron-glia interactions during neurodegeneration. J. Neurosci., 20, 7964–7971.

40. Romagnoli, M., Mineva, N.D., Polmear, M., Conrad, C., Srinivasan, S., Loussouarn, D., Barillé-Nion, S., Georgakoudi, I., Dagg, Á., McDermott, E.W. et al. (2014) ADAM8 expression in invasive breast cancer promotes tumor dissemination and metastasis. EMBO Mol. Med., 6, 278–294.

41. Knolle, M.D. and Owen, C.A. (2009) ADAM8: a new therapeutic target for asthma. Expert Opin. Ther. Targets, 13, 523–540.

42. Srinivasan, S., Romagnoli, M., Bohm, A. and Sonenshein, G.E. (2014) N-glycosylation regulates ADAM8 processing and activation. Journal of Biological Chemistry, 289, 33676–33688.

43. Hall, T., Leone, J.W., Wiese, J.F., Griggs, D.W., Pegg, L.E., Pauley, A.M., Tomasselli, A.G. and Zack, M.D. (2009) Autoactivation of human ADAM8: a novel pre-processing step is required for catalytic activity. Biosci Rep, 29, 217–228.

44. Hu Frisk, J., Pejler, G., Eriksson, S. and Wang, L. (2022) Structural and functional analysis of human thymidylate kinase isoforms. Nucleosides Nucleotides Nucleic Acids, 41, 321–332.

45. Liu, Y., Marks, K., Cowley, G.S., Carretero, J., Liu, Q., Nieland, T.J.F., Xu, C., Cohoon, T.J., Gao, P., Zhang, Y. et al. (2013) Metabolic and functional genomic studies identify deoxythymidylate kinase as a target in LKB1-mutant lung cancer. Cancer Discov., 3, 870–879.

46. Löffler, M., Carrey, E.A. and Zameitat, E. (2018) New perspectives on the roles of pyrimidines in the central nervous system. Nucleosides Nucleotides Nucleic Acids, 37, 290–306.

47. Crawford, D.C., Acuña, J.M. and Sherman, S.L. (2001) FMR1 and the fragile X syndrome: human genome epidemiology review. Genetics in medicine, 3, 359–371.

48. Santoro, M.R., Bray, S.M. and Warren, S.T. (2012) Molecular mechanisms of fragile X syndrome: a twenty-year perspective. Annu Rev Pathol, 7, 219–245.

49. Zafarullah, M., Tang, H.T., Durbin-Johnson, B., Fourie, E., Hessl, D., Rivera, S.M. and Tassone, F. (2020) FMR1 locus isoforms: potential biomarker candidates in fragile X-associated tremor/ataxia syndrome (FXTAS). Sci Rep, 10, 11099.

50. Sittler, A., Devys, D., Weber, C. and Mandel, J.-L. (1996) Alternative Splicing of Exon 14 Determines Nuclear or Cytoplasmic Localisation of FMR1 Protein Isoforms. Human Molecular Genetics, 5, 95–102.

51. Fu, X., Zheng, D., Liao, J., Li, Q., Lin, Y., Zhang, D., Yan, A. and Lan, F. (2015) Alternatively spliced products lacking exon 12 dominate the expression of fragile X mental retardation 1 gene in human tissues. Mol Med Rep, 12, 1957–1962.

52. Murillo-de-Ozores, A.R., Rodríguez-Gama, A., Carbajal-Contreras, H., Gamba, G. and Castañeda-Bueno, M. (2021) WNK4 kinase: from structure to physiology. Am. J. Physiol. Renal Physiol., 320, F378–F403.

53. Moriguchi, T., Urushiyama, S., Hisamoto, N., Iemura, S.-I., Uchida, S., Natsume, T., Matsumoto, K. and Shibuya, H. (2005) WNK1 regulates phosphorylation of cation-chloride-coupled cotransporters via the STE20-related kinases, SPAK and OSR1. J. Biol. Chem., 280, 42685–42693.

54. San-Cristobal, P., Ponce-Coria, J., Vázquez, N., Bobadilla, N.A. and Gamba, G. (2008) WNK3 and WNK4 amino-terminal domain defines their effect on the renal Na+-Cl-cotransporter. Am. J. Physiol. Renal Physiol., 295, F1199–1206.

