## Supplementary_information for "Uncovering the translatome impact of transcriptome induced diversity in eukaryotes: framework and innovative insights"

**S1: Maximum $FrSim_{exon}$ for $\delta_{exon}$ <0 of a pair of isoforms in a gene**

The distribution of $FrSim_{exon}$ computed for isoform pairs lies below a straight line (not shown in the Figure 3) for $\delta_{exon}$ <0. This line would represent the maximum possible of $FrSim_{exon}$ score for a given $\delta_{exon}$. Here, we describe the theoretical maximum value of $FrSim_{exon}$ score for a given $\delta_{exon}$. As mentioned in the main text, $FrSim_{exon}$ is the ratio of number of identical featured exon to the number of RISO exons of a gene, whereas the $\delta_{exon}$ is difference between the numbers of exons in mIDI to RISO.

$$max(FrSim_{exon}) = \frac{Nexon_{mIDI}}{Nexon_{RISO}} \qquad (eq\ 1)$$

$$x_{score} = 1 + \delta_{exon} \qquad (eq\ 2)$$

$$x_{score} = 1 + \frac{Nexon_{mIDI} - Nexon_{RISO}}{Nexon_{RISO}} = \frac{Nexon_{mIDI}}{Nexon_{RISO}} \qquad (eq\ 3)$$

The $max(FrSim_{exon})$ is be give by *eq 1*, as the maximum possible value of numerator will the number of exons in mIDI. If we perform algebraic manipulation (as given in *eq 2* and *eq 3*) of adding 1 to $\delta_{exon}$, the numerical values ($x_{score}$) determined from of *eq 1 and eq 3* are same. Thus, the $max(FrSim_{exon})$ is 1+ $\delta_{exon}$ when $\delta_{exon}$ < 0 will form a straight line in plot of $\delta_{exon}$ and $FrSim_{exon}$.

**S2: Deviation from expected distribution, chi square goodness of fit (Supplementing figure 4 and 5)**

To investigate whether prevalence of AS events in Figure 4 and 5 (in the main text) varies with isoform divergence, we examined consistency of these event distributions across various sID bins. With this assumption, we exploited $\chi 2$ goodness of fit by considering the sID bins as distinct categories and the expected AS events computed from observed occurrences. The AS events across categories (sID bins) does not follow the expected distribution ($\chi 2$ test is statistically significant). Subsequently, we derived the factor (bin), which has the most influence on $\chi 2$ test statistic by subdividing bins and serially eliminating one factor at a time to find the one having the most varying distribution from expected frequency [1] (Supplementary file SuppDataExcel.xlsx and spreadsheet F4_singleEvent_coding F4_singleEvent_noncoding and F5_doubleEventCoding).

**S3: Overview of ENACT Database**

Using our nomenclature, we have annotated exons of five widely studied model organisms, *viz. Caenorhabditis elegans, Drosophila melanogaster, Danio rerio, Mus musculus,* and *Homo sapiens* and documented them in the ENACT resource database (enactdb). Database is publicly available at http://www.iscbglab.in/enactdb/. The Table S1 summarizes the number of annotated exons/transcripts of genes encoded in five organisms available in enactdb.

**Table S1: Summary statistics of gene/transcript/exon in five model organisms.**

| Organism | Number of protein coding genes | Number of transcripts | Number of exons |
|---|---|---|---|
| *C. elegans* (Ce) | 19,972 | 28,534 | 1,25,054 |
| *D. melanogaster* (Dm) | 13,972 | 30,755 | 65,958 |
| *D. rerio* (Dr) | 26,374 | 48,821 | 2,68,035 |
| *M. musculus* (Mm) | 22,134 | 92,400 | 2,32,520 |
| *H. sapiens* (Hs) | 20,443 | 1,30,739 | 2,41,910 |

**Additional Files**

SuppDataExcel.xlsx

Worksheet named as "**F1_Identity_Coverage_gene_wise"** contains data concerning figure 1, where for five organisms, the taxonomic id is used to indicate organism (Taxonomic id are as follows: 6239: *C. elegans*; 7227: *D. melanogaster*, 7955: *D. rerio*; 10090: *M. musculus;* and 9606: *H. sapiens*). The isoform length (mIDI and RISO), identity, and coverage from RISO are listed per gene.

Worksheets named as "**F3_exon_composition_l3, F3_exon_composition_l2 and F3_exon_composition_l1**" contains data concerning figure 3, where data is binned per gene for exon composition between mIDI and RISO as function of identity and coverage. L3 to L1 in suffix in end of spreadsheets depicts the incorporation of blocks, where L3 is all blocks, L2 is 2 blocks (Block 2 and Block 1) and L1 is only Block 2.

Worksheet named as **"F4_singleEvent_coding F4_singleEvent_noncoding and F5_doubleEventCoding"** comprising supplement data to figure 4. The observed values for sID and expected (as inferred from unsegregated data) are listed, with $\chi^2$

goodness of fit calculations. Colored scheme and iterative procedure (mentioned also in spreadsheet) was used to recalculate statistic-associated p-value on removing sID bin with largest deviation in order to identify sID bins with contrast from expected distribution.

Database: ENACTdb (http://www.iscbglab.in/enactdb/) provided utility and usage of nomenclature is discussed in section "Overview of ENACT Database" and "Description of ENACT nomenclature" and illustrates AS event depiction and protein feature association in visually appealing manner.

References:


1. Zar JH: *Biostatistical analysis (5th edition), Chapter 22.* Pearson Education India; 2010.
